# Dysregulated actin dynamics and cofilin correlate with TDP-43 pathology in sporadic amyotrophic lateral sclerosis

**DOI:** 10.1101/2023.08.28.555209

**Authors:** Cyril Jones Jagaraj, Prachi Mehta, Sonam Parakh, Sina Shadfar, Shafi Jamali, Alexandra K Suchowerska, Jessica Sultana, Thomas Fath, Julie D Atkin

## Abstract

Amyotrophic lateral sclerosis (ALS) is a fatal, rapidly progressive neurodegenerative disorder affecting motor neurons, that overlaps significantly with frontotemporal dementia (FTD). Most cases are sporadic (90%) with undefined aetiology, but pathological forms of TAR-binding protein 43 (TDP-43), involving its misfolding, aggregation and mislocalisation from the nucleus to the cytoplasm, are present in motor neurons in almost all cases (97%) and ∼45% FTD cases. Actin is the most abundant protein in eukaryotic cells, with structural roles in the cytoskeleton and diverse signalling functions. This includes neuronal-specific roles in dendritic spines, synapses, axonal growth cones, and plasticity. Actin is in constant dynamic equilibrium between two forms: free monomeric, globular actin (G-actin) and polymeric, filamentous actin (F-actin). Actin dynamics is regulated by several key actin-binding proteins, including tropomyosin 4.2 (Tpm4.2) and cofilin, which depolymerises actin filaments. Cofilin is activated by phosphorylation at Ser3 via LIM domain kinase1/2 (LIMK1/2), which is also regulated by phosphorylation via Rac1/cdc42. Here we demonstrate that actin dynamics is closely associated with pathological TDP-43 in ALS. More F-actin relative to G-actin was detected in lumbar spinal cords from both sporadic ALS patients and a mouse model displaying TDP-43 pathology (rNLS), and in neuronal cells expressing cytoplasmic TDP-43. Hence actin dynamics is dysregulated in sporadic ALS, resulting in more actin polymerization. We also detected increased levels of Tpm 4.2, Rac1/cdc42, and increased phosphorylation of both LIMK1/2 and cofilin, in sporadic ALS patients. TDP-43 also physically interacted with actin *in vitro* and in cell lysates, providing additional insights into actin dysregulation in ALS. rNLS mice display motor neuron loss and key ALS/MND behavioural phenotypes, and increased cofilin phosphorylation was also detected in these animals at symptom onset, implying that actin dynamics actively contributes to neurodegeneration. Moreover, pharmacological induction of actin polymerization produced features typical of pathological TDP-43 (cytoplasmic mis-localisation and formation of inclusions and stress granules) implying that actin dysregulation contributes to TDP-43 pathology in ALS. Importantly, we also detected more cofilin phosphorylation in spinal motor neurons from sporadic patients compared to healthy controls, revealing that our observations are clinically relevant and present in the relevant cell type. This study therefore identifies dysregulated actin dynamics as a novel disease mechanism associated with TDP-43 pathology and hence most ALS cases. It also implies that regulating cofilin or LIMK1/2 phosphorylation may be a novel therapeutic strategy in ALS, FTD and other diseases involving TDP-43 pathology.

## Introduction

Amyotrophic lateral sclerosis (ALS) is a fatal, rapidly progressing neurodegenerative disorder affecting motor neurons, that overlaps clinically and pathologically with frontotemporal dementia (FTD).^1,2^ Almost all cases of ALS (97%) and ∼half % of FTD cases display pathological forms of TAR DNA-binding protein 43 (TDP-43) in affected tissues.^3,4^ TDP-43 is a RNA /DNA binding protein localised primarily in the nucleus, but pathological forms misfold, aggregate and mis-localise to the cytoplasm in ALS/FTD.^5–7^ TDP-43 pathology is strongly linked to toxicity and neurodegeneration,^8–10^ thus is central to ALS/FTD. During cellular stress, TDP-43 is also recruited to stress granules, heterogeneous transient structures composed of proteins, pre-mRNAs, and untranslated mRNA transcripts, that are increasingly associated with neurodegeneration.^8,11,12^

Actin is a major cytoskeletal protein and the most abundant protein in many eukaryotic cells (1-5% total), particularly muscle (10%).^13^ It is highly conserved, and its functions include vesicle/organelle movement, motility, maintenance of cell shape and muscle contraction.^14^ Actin exists in either monomeric, globular form (G-actin) or polymerised into short (5 nm diameter), linear filaments (F-actin) that generate movement and transport cargo using myosin motors.^15^ ‘Actin dynamics’ refers to the steady state assembly/disassembly of F-actin filaments, which is regulated by multiple actin-associated proteins, including tropomyosin and cofilin.^14,16^ Tropomyosin’s are a large group of proteins (40 isoforms) that co-polymersise with actin, thereby regulating its functional diversity.^17^ Tropomyosin also modulates the binding of actin-binding proteins to F-actin in an isoform-dependent manner.^18^ Cofilin has a dual function in dissembling F-actin filaments by both depolymerizing and severing actin.^19^ Phospho-regulation of cofilin on serine 3 is central to actin dynamics because phosphorylation inhibits its activity, whereas dephosphorylation activates its functions.^20^ Tropomyosin 4.2 (Tpm-4.2), which is highly enriched in the post-synaptic compartment,^21,22^ promotes cofilin phosphorylation and hence actin polymerisation.^23,24,25^ Cofilin is tightly regulated by LIMK1 and LIMK2 via RhoGTPases,^26^ master regulators of actin dynamics that switch between an inactive GDP bound form and an active GTP bound form.^27^ LIMKs are essential for spine formation, synaptic function, plasticity and memory formation, membrane trafficking and cell death.^19,28,29^

Actin dynamics and cofilin are linked to many processes associated with neurodegeneration,^30^ including redox sensing,^31,32^ apoptosis,^33,34^ DNA repair,^35^ cellular transport,^36^ and autophagy,^33,37^ lipid metabolism,^38^ maintenance of nuclear architecture,^39–41^ and aging.^34^ Actin also has neuronal-specific functions in dendritic spines, synapses and axonal growth cones and the G-actin: F-actin ratio is important in neuronal development and memory formation.^42–45^ Dysfunctional actin dynamics and/or cofilin abnormalities are well documented in Alzheimer’s,^46^ Parkinson’s,^47^ Huntington’s diseases,^48^ and spinal muscular atrophy.^49^ Whilst defects in other cytoskeletal components, particularly neurofilaments^50^ and microtubules,^51^ are described in ALS/FTD, the role of actin dynamics in pathogenesis remains unclear. However, mutations in profilin 1, which catalyses actin polymerisation, are present in familial ALS^52^ and actin dynamics is dysregulated in a small cohort of familial C9orf72 ALS patients^53^ and mutant superoxide dismutase (SOD1)^G93A^ mice at disease end stage.^54^

Here we show that actin dynamics is dysregulated by TDP-43 pathology. More F-actin and an increased F-actin: G-actin ratio were present in tissues from sporadic ALS (SALS) patients, and mouse (rNLS) and cellular models displaying TDP-43 pathology.^55^ More phosphorylation of cofilin and LIMK1/2, and increased Tpm-4.2 levels, were detected in SALS patients or rNLS mice prior to neurodegeneration. Moreover, pharmacological induction of actin polymerisation induced mis-localisation of TDP-43 to the cytoplasm and the formation of stress granules, implying that dysregulating actin dynamics induces TDP-43 pathology. These findings provide novel insights into the pathophysiology of TDP-43 in ALS/FTD.

## Materials and Methods

### Constructs

pLenti-C-monomeric (m) green fluorescent protein (GFP) (empty vector), pLenti-C-TDP-43-mGFP, and pLenti-C-TDP-43ΔNLS-mGFP were obtained from Dr. Adam Walker, University of Queensland. Codon-optimized human TDP-43 tagged with histidine (6X His) was sub-cloned into LIC-A plasmid (from Dr. Begona Heras, La Trobe University) with T7 promoter for bacterial expression.

### TDP-43 rNLS mice cortical lysate preparation

Mice were anesthetized with ketamine (100 mg/kg) and xylazine (10 mg/kg) and perfused intracardially with phosphate-buffered saline (PBS). The brain was removed, bisected through the midline, and immediately stored at -80^D^C. The cortices were lysed in 100mM ammonium bicarbonate, pH 7.6 containing 1% sodium deoxycholate with 1X protease/phosphatase inhibitor cocktail (Roche, #05892970001). Samples were then sonicated, spun at 4^°^C ∼100,000*g* for 30 min and the supernatant stored at -80^°^C. Total protein concentration was quantified (Pierce™ BCA Protein Assay Kit), following the manufacturer’s instructions.

### Human spinal cord lysate preparation

Human lumbar spinal cord segments (L3–L5) from nine MND patients and seven control individuals without neurological/psychiatric disease, were provided by the Victorian Brain Bank (**Table 1**). 100mg of tissue was lysed in radioimmunoprecipitation assay buffer (RIPA, 50mm Tris–HCl, pH 7.5, 150mm NaCl, 0.1% (w/v) SDS, 1% (w/v) sodium deoxycholate and 1% (v/v) Triton X-100) with 1% (v/v) protease inhibitor cocktail (Roche). After homogenization, samples were centrifuged for 10 min at 100,000g, and the supernatant frozen at −80^°^C. Protein lysates were quantified using a BCA assay following the manufacturer’s instructions.

### Insoluble cellular fraction preparation

NSC-34 cells expressing either EGFP-tagged TDP-43, EGFP only, or EGFP-tagged TDP-43 ΔNLS, or untransfected cells, were lysed with RIPA buffer with protease and phosphatase inhibitors (Sigma #4693159001) and incubated on ice for 20 min. Following centrifugation at 100,000xg for 30 min at 4^°^C the RIPA-soluble fraction (supernatant) was discarded. The RIPA-insoluble fraction (pellet) was washed twice with RIPA buffer to remove residual soluble proteins and 50ul of RIPA buffer containing 2% SDS was added followed by sonication and centrifugation at 100,000g at 22^°^C for 30 min. The supernatants were retained as the insoluble protein fractions.

### G-actin/F-actin *in vivo* assays

#### Cell lysates

F- and G-actin fractions from lysates were prepared using a G-Actin/F-Actin *in Vivo* Assay Biochem kit (Cytoskeleton, #BK037) following the manufacturer’s instructions. Briefly, transfected cells were washed in PBS and incubated in 200μl of lysis buffer (LAS2) at 37^°^C for 10 minutes. Lysates (100μl) were centrifuged at 500 g for 5 min at 37^°^C to remove debris, and the supernatant was centrifuged at ∼100,000 g for 1hr at 37^°^C. The G-actin rich supernatant and F-actin rich pellet were collected: 100μl of F-actin depolymerizing buffer was added to the latter followed by incubation on ice for 1 hr. The G-actin: F-actin ratio was quantified by Western blotting.

#### Mouse cortical tissues

TDP-43 rNLS mice and littermate controls were deeply anesthetized using ketamine/xylazine and intracardially perfused with ∼ 20 ml 0.9% saline solution. Cortices were dissected, frozen on dry ice and stored at -80^°^C until required when 50μg was lysed in 100 *µ*l LAS2 buffer. F/G-actin rich fractions were prepared as above.

#### Human sporadic ALS patient spinal cords

To 100mg each tissue (**Table 1**), 1ml of LAS2 buffer was added and F/G-actin fractions were prepared as above.

### Recombinant TDP-43 protein purification

Codon-optimized human TDP-43-His LIC-A constructs were transformed into T7 express (New England Biolabs, #C3029J) *E. coli*, which constitutively expresses the chaperone DsbC to promote cytoplasmic disulfide bond formation and hence recombinant protein folding/expression. *E. coli* were grown in 2L LB medium containing 100 μg/ml ampicillin at 30^°^C until OD_600_ = 0.6. TDP-43 expression was induced by addition of 1mM isopropyl β-D-thiogalactopyranoside (Sigma, #11411446001) for either 4 hr at 30^°^C or 24 hr at 18^°^C, and cells harvested by centrifugation and stored at -80^°^C. Recombinant human TDP-43 was expressed as insoluble cellular inclusion bodies. Pellets were resuspended in lysis buffer (20mM Tris-HCL pH 8.0, 300mM NaCl; 4ml/100ml culture) with freshly added protease inhibitors (Roche #11697498001: 1 tablet/50ml) and lysozyme (0.1mg/ml), followed by incubation at room temperature (RT) on a rotary mixer for 15mins. Following sonication (10 cycles of 20s on /5s off, at 40-50% power), 10 *µ*g/ml DNase was added followed by incubation with mixing for 15 min and centrifugation at 20,000 g at 4^°^C for 30 min. Pellets were resuspended in solubilization buffer (6 M guanidinium hydrochloride, 20mM Tris-HCL pH 8.0, 300mM NaCl, 20mM imidazole; 5ml/100ml culture), incubated at RT on a rotary mixer for 2+ hours, and centrifuged at 20,000 g for 30 min at 4^°^C. The supernatant containing recombinant TDP-43 was purified by affinity chromatography using Ni-NTA columns pre-equilibrated with solubilization buffer. Supernatant was applied, then 10ml of wash buffer (4M guanidinium hydrochloride, 20mM Tris-HCl pH 8.0, 300mM NaCl, 20 mM imidazole) was added at decreasing GuHCL concentrations (4M>2M>1M>0.5M>0M), to refold denatured TDP-43 on the column. Finally, recombinant TDP-43 was eluted with elution buffer (20mM Tris-HCl pH 8.0, 300mM NaCl, 250 mM imidazole) and stored at -80^°^C. Purity was assessed by SDS/PAGE using Coomassie Brilliant Blue staining and concentration determined spectrophotometrically at absorbance of 280 nm. Stability was confirmed after incubating ∼0.1-1*µ*g at RT for 4 hr followed by SDS/PAGE and Coomassie staining, detected by lack of cleaved/degraded TDP-43.

### Immunoprecipitation

∼30µg of purified recombinant TDP-43 and 10*µ*g of rabbit skeletal muscle actin (Cytoskeleton, #BK037) were incubated with Ni-NTA beads at 4^°^C overnight on a rotating wheel, followed by centrifugation at 2000 rpm for 5 min. The pellet was washed twice with wash buffer (10mM Tris/Cl pH 7.5, 150mM NaCl, 0.5mM EDTA). The immunoprecipitate was released by boiling with 1xSDS sample loading buffer for 5 min, followed by centrifugation at 15,000 g for 2 min. GFP-Trap immunoprecipitation assay was performed following the manufacturer’s instructions (ChromoTek, #gta).

### Immunohistochemistry

Human thoracic spinal cord segments (L3-L5) from three SALS patients (clinical diagnosis confirmed post-mortem, P7-9), and two controls without evidence of neurological/psychiatric disease (C6-7), were provided by the Victorian Brain Bank **(Table 1)**. Sections were pre-heated at 70^°^C for 30 min, and paraffin was removed by washing; 2x xylene, 2x 100% ethanol, 1x 95% ethanol, 1x 70% ethanol and 2x distilled water. Antigen retrieval was performed by boiling sections in 10mM sodium citrate buffer for 30 min followed by blocking with 5% goat serum containing 0.1% Tween 20 in PBS for 1 hr. Primary antibodies, rabbit anti-phospho-cofilin (CST, #3311, 1:100) and mouse anti-SMI-32 (BioLegend, # SMI-32R, 1:250), were applied in blocking buffer at 4^°^C overnight, followed by secondary antibodies; donkey anti-rabbit Alexa Fluor 594 (Invitrogen, #A-21207, 1:200) and rabbit anti-mouse Alexa Fluor 488 (Invitrogen, #A-11059, 1:200) for 1hr in the dark at RT. After washing with PBS, staining of the nuclei was performed using Hoechst stain 33258 (Sigma, #94403). Sections were imaged using 63×oil objective lens and Zeiss LSM 880 inverted confocal laser-scanning microscope. Quantification was performed from 13 neurons each in both controls and SALS patients.

### Immunocytochemistry

NSC-34 cells cultured on 13mm coverslips (Menzel-Glasser) in Dulbecco’s Modified Eagle Medium media (DMEM, Gibco) were transfected with TDP-43 or control plasmids using Lipofectamine 2000 (Invitrogen) for 48hr following manufacturer’s instructions. To induce actin polymerization, cells were treated with 1%DMSO or 1µM Jasplakinolide in DMSO (Sigma, #J4580) for 1hr, then washed in PBS and fixed in 4% PFA for 10 min, permeabilized with 0.1% Triton X-100 in PBS for 5 mins and blocked with 5%BSA in PBS for 30 mins. Mouse anti-HuR (Invitrogen, #390600, 1:100) primary antibody in blocking buffer was incubated with cells at 4^°^C overnight and secondary antibody, donkey anti-rabbit Alexa Flour 594 (Invitrogen, #A21207, 1:200) was added for 1 hr. After three washes in PBS, cells were treated with 0.5 *µ*g/ml Hoechst 33258 (Sigma, #94403) and coverslips mounted onto slides in fluorescent mounting media (Dako). Mislocalisation of TDP-43 was analyzed from a minimum of 100 cells/group from three independent experiments.

### SDS-PAGE and Western blotting

Samples (20µg protein) were subjected to either 4-15% SDS-PAGE (Bio-Rad), or 10% SDS-PAGE for Tpm-4.2, and transferred onto nitrocellulose membranes according to manufacturer’s instructions (Bio-Rad). Blots were pre-incubated in blocking solution containing 5%(w/v) skim milk in Tris-buffered saline (TBS), followed by incubation overnight at 4°C in primary antibodies diluted in 5% (w/v) BSA in Tris buffered saline: anti-cofilin 1G6A2 (Proteintech, 66057-I-Ig, 1:3000), anti-phospho-cofilin (Ser3) (CST, #3311, 1:1000), anti-GAPDH (Proteintech, #60004-1-Ig, 1:5000), anti-TDP-43 (Cosmo Biotech, #TIP-TD-P09, 1:1000), anti-GFP (Abcam, #ab290, 1:2000), anti-Rac1/cdc42 (CST, #4651, 1:1000), anti-phospho-LIMK1 (Thr 505,507)(CST, #3841, 1:500), anti-LIMK1 (CST, #3842, 1:1000), anti-β-actin (Sigma, #A2228, 1:4000), anti-TDP-43 (Proteintech, #10782-2-AP, 1:1000), anti-β-actin clone C4 (Merck, #MAB1501, 1:1000) and anti-Tpm-4.2 (Suchowerska et al,^22^ 1:1000). Immunoreactivity was revealed using the Clarity™ ECL Western Blotting Substrate kit (BioRad) and images obtained using ChemiDoc MP with Image Lab™ software (BioRad). The intensity of each band relative to GAPDH was quantified using ImageJ (v. 1.47; National Institutes of Health).

### Confocal Microscopy

Cells were photographed with 63x/na=1.4 or 100x/na=1.46 objectives on a Zeiss LSM 880 inverted confocal laser-scanning microscope. Photomultiplier sensitivities and offsets were set to a level in multichannel imaging, at which bleed through effects from one channel to another were absent. All confocal images were obtained under equivalent conditions of laser power, pinhole size, gain, and offset settings between the groups. All laser exposures were kept constant between groups.

### Statistics

Human post-mortem sample size was based on tissue availability. Data are presented as mean value with standard deviation (SD). Statistical comparisons between group means were performed using GraphPad Prism 8.02 software (Graph Pad, Inc.). A t-test or a one-way ANOVA, followed by a post hoc Tukey test for multiple comparisons, was used when justified. The significance threshold was set at *p* = 0.05. The number of independent repeats (*n*), statistical tests used, and statistical significances (*p* values) are specified in each figure legend. Immunofluorescence and immunohistochemistry experiments were performed blinded.

### Data availability

The data supporting the results of this study are available upon reasonable request to the corresponding author.

## Results

### Actin dynamics is perturbed in sporadic ALS patients

First, actin dynamics was investigated in lumbar spinal cord tissues from SALS patients and non-neurological controls, using a G-actin and F-actin *in vivo* assay, followed by western blotting for β-actin. This assay uses a specific detergent-based lysis buffer to isolate G-actin and F-actin fractions. Quantification revealed a significant decrease in the levels of G-actin in human SALS lysates compared to controls (0.30-fold, *\*p*=0.0197) (**Fig 1A-B**), demonstrating that less monomeric actin is present. Correspondingly, significantly more F-actin was present in SALS tissues (2.51-fold, *\*p*=0.0474) (**Fig 1C-D**), revealing more polymerised actin. Furthermore, the G: F-actin ratio was significantly decreased in SALS compared to control patients (0.13-fold, *\*\*p*=0.0016) (**Fig 1E**), together implying that actin dynamics is disturbed in SALS lumbar spinal cords.

**Figure 1.**
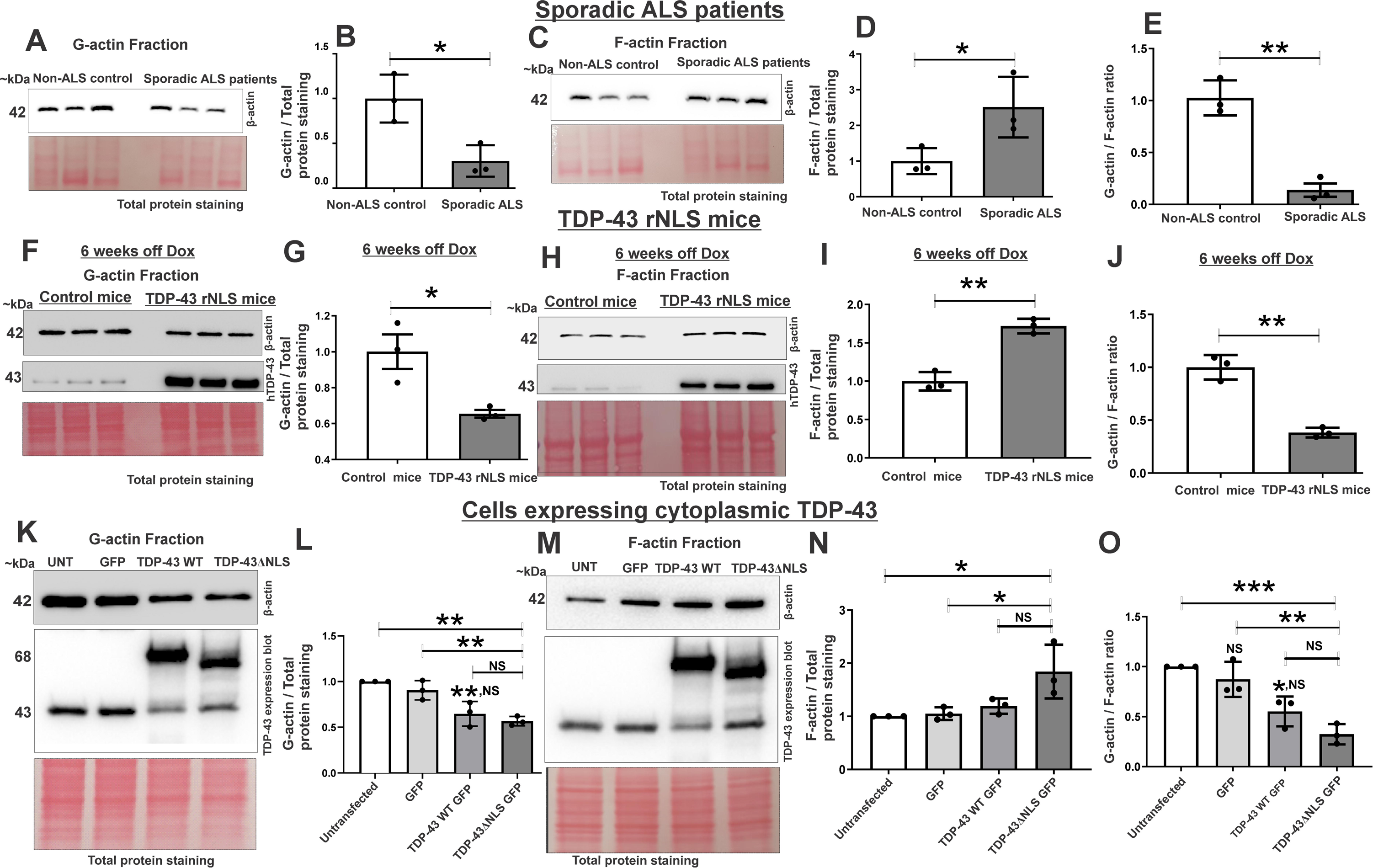
Actin dynamics is disturbed in sporadic ALS patients, TDP-43 rNLS mice and in cells expressing cytoplasmic TDP-43. **Sporadic human ALS tissues (A and C)** Western blotting of G-actin and F-actin fractions of lumbar spinal cord tissues from non-neurological controls and SALS patients using an anti-β-actin antibody. Total protein staining using Ponceau S was used as a loading control. Approximate molecular weight markers in kDa are shown on the left. **(B)** Quantification of the G-actin fractions of the blots in (A). Relative intensity of G-actin to total protein in fractions of control and SALS patients. Data represented as mean ± SD (*n* = 3), unpaired t-test, two-tailed *\*p*<0.05. A significant decrease in the levels of G-actin in SALS patients compared to controls was detected. (D) Quantification of F-actin fractions of blots shown in **(C)**. Relative intensity of F-actin to total protein in fractions, mean ± SD (*n* = 3), unpaired t-test, two-tailed *\*p*<0.05. (E) Quantification of G-actin/F-actin ratio from (B) and **(D)**, mean ± SD (*n* = 3), unpaired t-test, two-tailed *\*\*p*<0.01. A significant decrease in the G-actin: F-actin ratio in SALS patients compared to control patients was detected. **TDP-43 rNLS mice (F and H)** Western blotting of G-actin and F-actin fractions prepared from TDP-43 rNLS mice and littermate controls at 6 weeks off Dox (late symptomatic stage) using anti-β-actin and anti-TDP-43 antibodies. Total protein staining with Ponceau S was used as a loading control. TDP-43 blot confirms transgene expression (hTDP-43 ΔNLS) in TDP-43 rNLS mice. Approximate molecular weight markers in kDa are shown on left. **(G)** Quantification of the G-actin fractions of western blots in (F). Relative intensity of G-actin to total protein staining in fraction of control mice and TDP-43 rNLS mice at 6 weeks off Dox. Data represented as mean ± SD (*n* = 3), unpaired t-test, two-tailed *\*p*<0.05. Significantly less G-actin is present in TDP-43 rNLS mice relative to control animals. **(I)** Quantification of the F-actin fractions of blots in (H). Relative intensity of F-actin to total protein staining fractions of control mice and TDP-43 rNLS mice 6 weeks off Dox. Graph represents average of at least two technical replicates of each lysate, mean ± SD (*n* = 3 mice), unpaired t-test, two-tailed *\*\* p*<0.01. Significantly more F-actin was present in TDP-43 rNLS mice relative to control animals. **(J)** Quantification of G-actin/F-actin ratio from (G) and (I), mean ± SD (*n* = 3), unpaired t-test, two-tailed *\*\*p*<0.01. A significant decrease in the G/ F actin ratio is present in 6 weeks off Dox TDP-43 rNLS mice compared to control mice. **NSC-34 cells (K and M)** Western blotting of G-actin and F-actin rich-fractions prepared from untransfected cells (UNT) and cells expressing empty vector, GFP only (GFP), wild-type TDP-43 or TDP-43 ΔNLS, using anti-β-actin and anti-TDP-43 antibodies. Total protein staining with Ponceau S was used as a loading control. TDP-43 blot confirms overexpression of TDP-43 WT and TDP-43-ΔNLS. Approximate molecular weight markers in kDa are shown on left. **(L)** Quantification of G-actin fractions from blots in (K). Relative intensity of G-actin to total protein in fractions. Data are represented as mean ± SD (*n* = 3), one–way Anova, Tukey’s multiple comparison \**p*<0.05, \*\**p*<0.01, *NS*, non-significant **(N)** Quantification of F-actin-fractions of blots in (M), mean ± SD (*n* = 3), one–way Anova, Tukey’s multiple comparison analysis \**p*<0.05, \*\**p*<0.01, *NS*, non-significant. **(O)** Quantification of G-actin: F-actin ratio from (L) and (N), mean ± SD *(n* = 3), one–way Anova, Tukey’s multiple comparison analysis \**p*<0.05, \*\**p*<0.01, \*\*\**p*<0.001, *NS*, non-significant.

### Actin dynamics is disturbed in TDP-43 rNLS mice

Next, actin dynamics was examined in a mouse model displaying TDP-43 pathology, rNLS^55^ . This model involves doxycycline (Dox)-suppressible expression of human TDP-43 with a mutated nuclear localization signal, driven by the neurofilament heavy chain promoter. Cytoplasmic TDP-43 accumulates in the cortex of these animals one week after human TDP-43 expression is initiated. Loss of cortical/spinal motor neurons, motor deficits, progressive neurodegeneration, and early death then follows.^55^ Cortical lysates from rNLS mice at six weeks off Dox (late symptomatic disease stage) (Supplementary Fig 1H), and from non-transgenic, littermate monogenic controls, were subjected to a G-actin/F-actin assay and western blotting for β-actin. A significant decrease in G-actin levels in rNLS mice compared to controls (0.65-fold, *\*p*=0.0274) (**Fig 1F-G**) revealed that less monomeric actin was present. Correspondingly, more F-actin, representing polymerised actin, was present in TDP-43 rNLS cortices compared to controls (1.71-fold, *\*\*p*=0.0012) (**Fig 1H-I**), and the G/F-actin ratio was also significantly reduced (0.38-fold, *\*\*p*=0.0010) (**Fig 1J**). Therefore, actin dynamics is disturbed in the cortex of TDP-43 rNLS mice.

### Overexpression of cytoplasmic TDP-43 disturbs actin dynamics

Next, GFP-only, GFP-tagged wildtype (WT) TDP-43, or TDP-43 with its nuclear localisation signal (NLS) removed (ΔNLS), were expressed in motor neuronal NSC-34 cells for 72h.^57^ Microscopy confirmed that ΔNLS TDP-43 was expressed mostly in the cytoplasm only (80% cells) or both nucleus and cytoplasm (20% cells), whereas WT TDP-43 was expressed in the nucleus only (90% cells) or in both the nucleus and cytoplasm (10% cells) (Supplementary Fig 1A). Actin dynamics was then examined in cell lysates using a G-actin and F-actin assay followed by western blotting for β-actin. Quantification revealed significantly less G-actin in cells expressing TDP-43-ΔNLS compared to untransfected (0.56-fold, *\*\*p*=0.0016) and GFP only (0.62-fold, *\*\*p*=0.0073), but not WT TDP-43, cells (**Fig 1K-L**). Similarly, significantly more F-actin was present in TDP-43-ΔNLS compared to untransfected (1.84-fold, *\*p*=0.0210) and GFP only, but not WT TDP-43, cells (1.75-fold, *\*p*=0.0287) (**Fig. 1M-N**). Whilst not statistically significant, trends of more F-actin and less G-actin in TDP-43-ΔNLS compared to TDP-43 WT cells were evident. There were no differences between GFP only and untransfected cells, demonstrating that actin dynamics was not perturbed by non-specific protein overexpression. Significantly less G-actin, but not F-actin, was present in WT TDP-43 (0.65-fold, *\*\*p*=0.0095), compared to untransfected, cells. The G/F-actin ratio was also reduced in TDP-43-ΔNLS and WT TDP-43 (0.55-fold, *\*p*=0.01) groups compared to untransfected cells (0.32-fold, *\*\*\*p*=0.0008) (**Fig 1O**), but only TDP-43-ΔNLS was different to GFP only cells (0.56-fold, *\*\*p*=0.0031) (**Fig 1O**). Given that either a major (ΔNLS) or minor (WT) proportion of TDP-43 was expressed in the cytoplasm in each case, these data imply that cytoplasmic TDP-43 dysregulates actin dynamics in ALS, resulting in more polymerised actin. Hence these results link TDP-43 pathology to actin in ALS.

### Actin polymerization induces features of TDP-43 pathology

We next examined whether perturbed actin dynamics initiates TDP-43 pathology by inducing the formation of F-actin pharmacologically. Jasplakinolide is a widely used, potent inducer of actin nucleation that stimulates actin polymerisation^58,59^. NSC-34 cells expressing WT TDP-43 GFP for 48 hr were treated with either 1µM Jasplakinolide or DMSO control for one hr. A G-actin and F-actin assay confirmed that actin polymerisation was induced by Jasplakinolide (Supplementary Fig 2A). No obvious signs of toxicity at this concentration were evident via light microscopy. Using fluorescent microscopy, TDP-43 mislocalisation and inclusion formation was next examined. Cells were classified as mislocalised when cytoplasmic TDP-43 was present (cytoplasm only, or both cytoplasm/nucleus), in contrast to nuclear TDP-43 (nucleus only). Significantly more cells expressed cytoplasmic TDP-43 following Jasplakinolide treatment compared to DMSO (\*\**p*=0.0014, **Fig 2A-B**). TDP-43 inclusions were identified by the presence of green-fluorescent, compact, cytoplasmic TDP-43 aggregates. Quantification revealed significantly more cells with TDP-43 inclusions following Jasplakinolide treatment compared to DMSO (\*\**p*=0.0043, **Fig 2A**, 2C). Hence stimulating actin polymerisation by Jasplakinolide induces mislocalisation and aggregation of TDP-43 - features of TDP-43 pathology.

**Figure 2.**
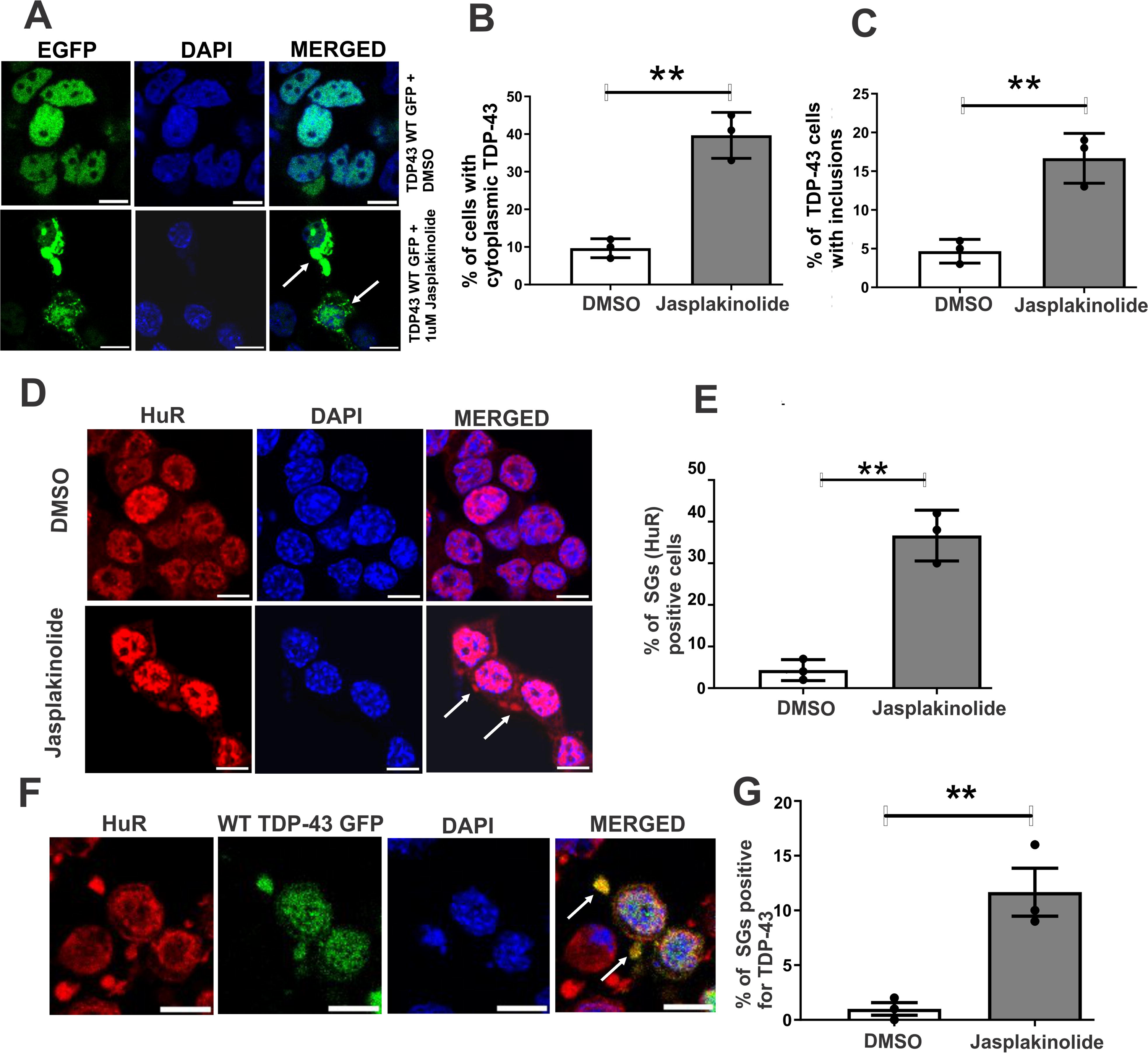
Actin polymerisation induces features of TDP-43 pathology in neuronal cells. **(A)** Fluorescent confocal microscopy of GFP-tagged WT TDP-43 expressing NSC-34 cells, with Hoechst staining. White arrows represent cytoplasmic TDP-43 inclusion-like structures. Scale bar = 10 μm. **(B)** Quantification of transfected cells in (A) displaying cytoplasmically localised GFP-tagged TDP-43 after treatment with DMSO or Jasplakinolide. >100 cells per group were examined. Data are represented as mean ±SD, *n*=3, unpaired t-test \*\**p*<0.01. **(C)** Quantification of transfected cells in (A) displaying cytoplasmic GFP-tagged TDP-43 inclusions after treatment with DMSO or Jasplakinolide. Data are represented as mean ± SD, *n*=3, unpaired t-test \*\**p*<0.01. **(D)** Fluorescent microscopy following HuR immunocytochemistry and Hoechst staining of TDP-43 WT GFP expressing NSC-34 cells (100+ cells per group) (*n*=3) treated with 1µM Jasplakinolide and DMSO. White arrows indicate HuR-positive stress granules. **(E)** Quantification of cells in (D) positive for stress granule marker HuR after treatment with Jasplakinolide or DMSO. Data are represented as mean ± SD, *n*=3, unpaired t-test \*\**p*<0.01. **(F)** Fluorescent confocal microscopy following HuR immunocytochemistry and Hoechst staining of TDP-43 WT GFP expressing cells. White arrows indicate TDP-43-positive stress granules, in which TDP-43 and HuR are co-localised. **(G)** Quantification of % of structures in (F) positive for both stress granule marker HuR and TDP-43 after treatment with Jasplakinolide or DMSO. Data are represented as mean ± SD, *n*=3, unpaired t-test \*\**p*<0.01.

Stress granules (SGs) are associated with pathological aggregation in ALS^60,61^, hence their formation was next investigated following 1µM Jasplakinolide treatment using immunocytochemistry for marker HuR. Fluorescent microscopy (**Fig 2D**) revealed the presence of HuR-positive cytoplasmic structures, and quantification revealed significantly more Jasplakinolide-treated cells contained SGs (*\*\*p*=0.0011, **Fig 2E**) compared to DMSO-treated cells. Hence, actin polymerisation induces SG formation (**Fig 2D-E**). Quantification of cells with co-localisation of TDP-43 and HuR (**Fig 2F**) revealed significantly more recruitment of TDP-43 to SGs following Jasplakinolide treatment than DMSO (*\*\*p*=0.0092, **Fig 2G**). Hence stimulating actin polymerisation with Jasplakinolide induces both SG formation and recruitment of TDP-43, providing further evidence of a link between actin polymerisation and TDP-43 pathology.

### TDP-43 immunoprecipitates with β-actin

We hypothesised that TDP-43 might interact directly with actin, which subsequently perturbs actin dynamics in ALS. GFP-tagged WT TDP-43, or control GFP only, were expressed in NSC-34 cells for 48 hours. Immunoprecipitation with anti–GFP coated magnetic beads followed by Western blotting revealed that significantly more actin was precipitated from TDP-43 lysates compared to untransfected (*\*p*=0.0103) and GFP only cells (*\*p*=0.0154) (**Fig 3A-B**). Western blotting using anti-GFP antibodies confirmed that TDP-43 was immunoprecipitated (**Fig 3A**). These results imply that TDP-43 interacts with actin.

**Figure 3.**
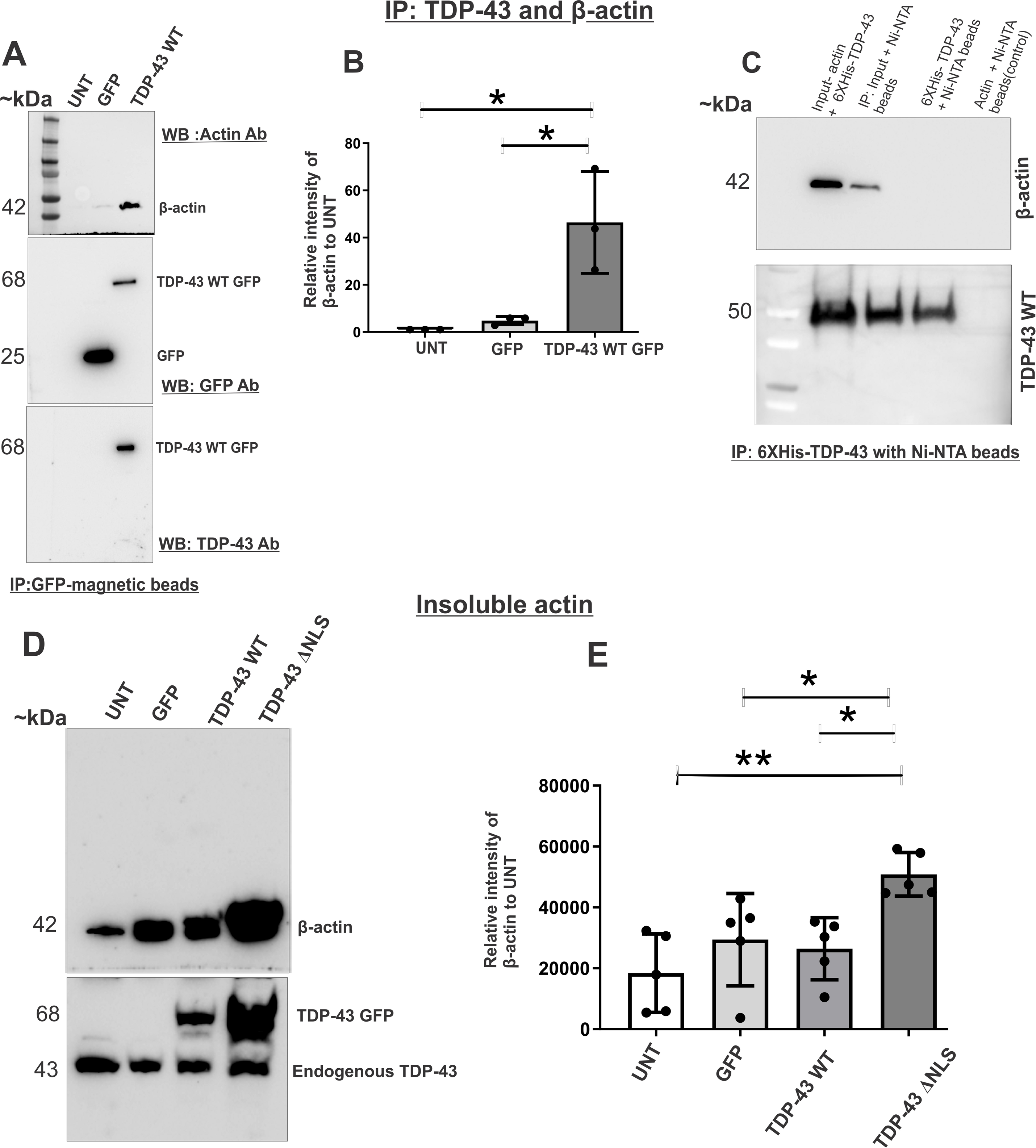
TDP-43 immunoprecipitates with β-actin and is enriched in insoluble fractions of neuronal cells expressing cytoplasmic TDP-43. **(A)** Western blotting using anti-β-actin, anti-GFP and anti-TDP-43 antibodies of NSC-34 lysates co-immunoprecipitated using GFP-magnetic beads prepared from untransfected cells (UNT), cells expressing empty vector (GFP) or GFP-tagged wild-type TDP-43. Approximate molecular weight markers are shown on the left. **(B)** Quantification of actin intensity from the blots shown in (A). Data are represented as mean ± SD (*n* = 3), one–way ANOVA followed by Tukey’s multiple comparison test: \**p*<0.05. **(C)** Precipitation of purified recombinant 6xHis-tagged TDP-43 and rabbit skeletal actin using Ni-NTA beads, followed by western blotting with anti-TDP43 and anti-β-actin antibodies. *Top panel* β -actin is precipitated by Ni-NTA, hence by TDP-43. *Bottom panel* Blot confirms that TDP-43 is present in the precipitant. Approximate molecular weight markers are shown on the left. **(D)** *Top panel* Western blotting of insoluble fractions using β-actin and TDP-43 antibodies from untransfected cells (UNT) and cells expressing empty vector (GFP), wild-type TDP-43-GFP or TDP-43ΔNLS-GFP. *Bottom panel* TDP-43 blot illustrates overexpression of GFP-tagged TDP-43 WT and TDP-43ΔNLS proteins. Approximate molecular weight markers are shown on the left. **(E)** Quantification of Western Blot replicates in (E). The graph depicts the absolute average intensities of β-actin in the insoluble fractions from untransfected NSC-34 cells and cells expressing GFP, WT TDP-43-GFP and TDP-43ΔNLS-GFP. Data are represented as mean ± SD, *n* = 5 independent experiments, One-way Anova, Tukey’s multiple comparison analysis \**p*<0.05, \*\**p*<0.01. More insoluble β-actin is present in cells expressing TDP-43 NLS compared to the other groups.

### TDP-43 directly precipitates with G-actin

The interaction between TDP-43 and actin was examined more specifically using purified proteins. Rabbit G-actin was purchased commercially, and recombinant TDP-43 was purified from *E. coli* expressing codon-optimised 6xHis tagged human WT TDP-43. SDS-PAGE of the eluted fractions, followed by staining with Coomassie Brilliant Blue (Supplementary Fig 3B), revealed that purified protein (<50kDa, as expected for 6xHis-tagged TDP-43: 0.8kDa + 43kDa), was present. Stability of the purified TDP-43 was assessed by SDS-PAGE followed by Coomassie staining, revealing that it was stable at RT for 4h (Supplementary Fig 3B). To examine the interaction between purified G-actin and recombinant TDP-43, precipitation with Ni-NTA agarose beads to pull-down TDP-43 was performed. Western blotting revealed that actin was precipitated with TDP-43 (**Fig 3C**) consistent with the cellular findings (**Fig 3A-B**), implying that these two proteins physically interact.

### Actin is enriched in insoluble fractions of cells expressing cytoplasmic TDP-43

Given that actin polymerises more in ALS and that actin co-precipitates with TDP-43, next it was examined whether actin is enriched in insoluble cellular fractions, implying that it is misfolded or aggregated in ALS. Therefore, in NSC-34 cells transfected as above for 72 hr, insoluble cellular fractions were subjected to Western blotting for β-actin and TDP-43. Quantification revealed significantly more insoluble actin was present in cells expressing ΔNLS TDP-43-GFP compared to untransfected (\**\*p*=0.0024), GFP-only (*\*p*=0.0476) and WT TDP-43-GFP cells (*\*p*=0.0217) **(Fig 3D-E)**. These data reveal that cytoplasmic TDP-43 is associated with more insoluble actin.

### Rac1/cdc42 and LIMK1 phosphorylation are enhanced in sporadic ALS patients

Rac1/cdc42 and RhoA are RhoGTPases that regulate actin dynamics via LIMK1/2 phosphorylation. Hence, their expression was next investigated in SALS patient lumbar spinal cords and non-neurological controls (**Table 1**) by Western blotting. Significantly more Rac1/cdc42 (4.3-fold, *\*p*=0.024) **(Fig 4A-B)** and phosphorylated LIMK1/2 (6.74-fold, \**p*=0.032) **(Fig 4C-D)** were present in SALS patients compared to controls. There was no change in total LIMK1 levels (0.942-fold, *ns p*=0.89) or loading (GAPDH) **(Fig 4E),** confirming that LIMK1 phosphorylation was specifically increased. Hence, more Rac1/cdc42 correlates with enhanced phosphorylation of LIMK1/2 in SALS patients.

**Figure 4.**
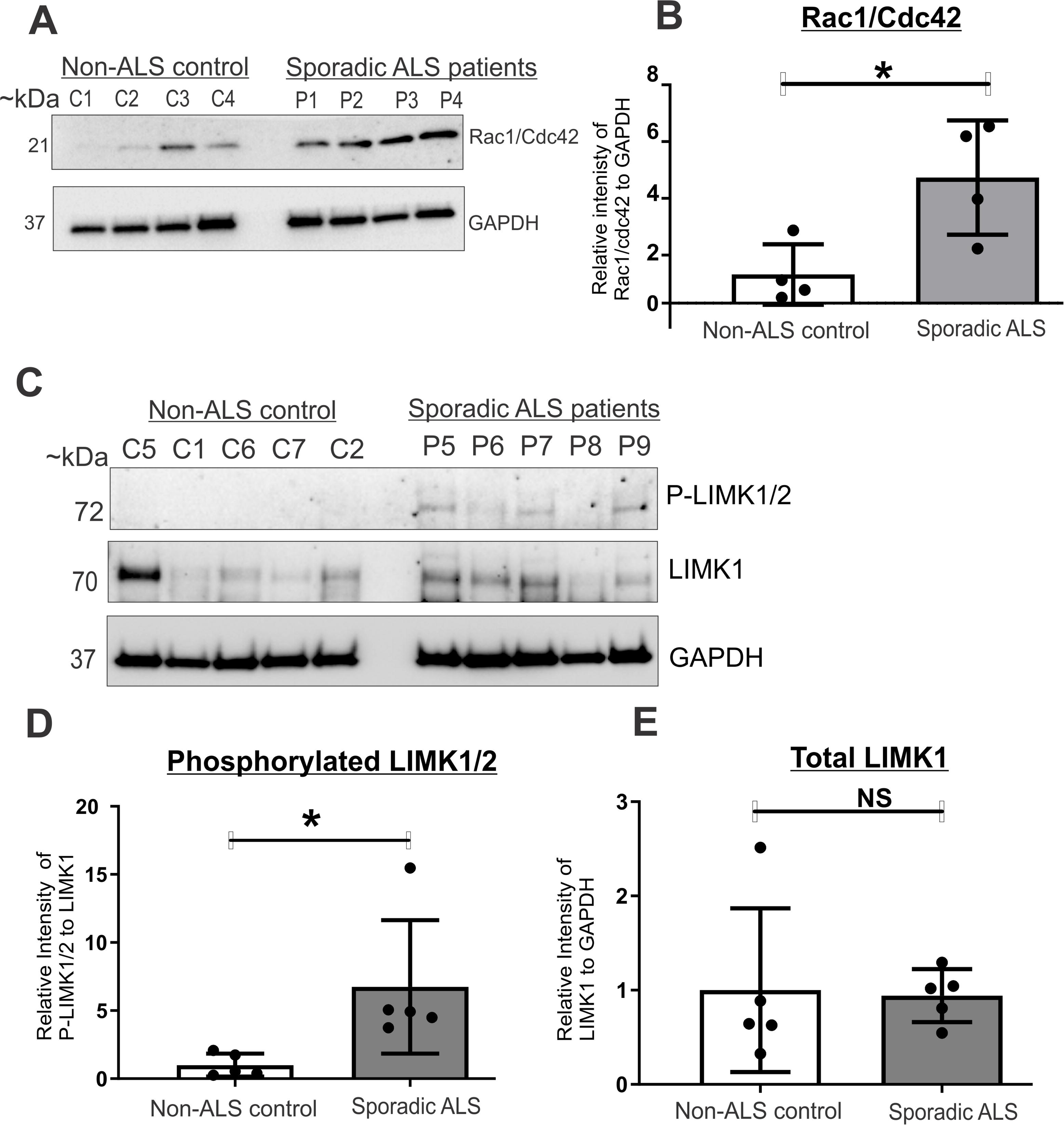
Phosphorylation of LIMK1 and Rac1/cdc42 is enhanced in sporadic ALS patient spinal cords. **(A)** Western blotting using anti-Rac1/cdc42 and anti-GAPDH antibodies in lumbar spinal cord lysates prepared from human SALS patients and non-ALS controls (*n*=4 per group). Approximate molecular weights (in kDa) are shown on the left. GAPDH was used as a loading control. **(B)** Quantification of relative intensity of Rac1/cdc42 to GAPDH from the western blots shown in (A) of SALS, normalized to non-ALS controls. Data are represented as mean ± SD (*n* = 4), unpaired t-test, two-tailed *\*p*<0.05. Significantly more Rac1/cdc42 is present in human sporadic ALS patients compared to non-ALS controls. **(C)** Western blotting using anti-phospho-LIMK1(Thr508)/LIMK2(Thr505), anti-LIMK1 and anti-GAPDH antibodies in spinal cord lysates prepared from human SALS patients and non-ALS controls (*n*=5). Approximate molecular weight (in kDa) is shown on the left. GAPDH was used as a loading control. Blots were reprobed using the same membrane, so loading is equivalent in each case. (**D)** Quantification of western blots shown in (C). The graph represents the relative intensity of phosphorylated LIMK1/2 to total LIMK1 in spinal cords of SALS patients, normalized to non-ALS controls. Data are represented as mean ± SD *(n* = 5), unpaired t-test, two-tailed *\*p*<0.05. Significantly more phosphorylated LIMK1/2 was present in human lumbar sporadic ALS patients compared to non-ALS controls. **(E)** Quantification of total LIMK1 to GAPDH ratio in western blots shown in (A). No significant difference in the levels of LIMK1 relative to GAPDH compared to controls was observed, *NS*, non-significant.

### Cofilin phosphorylation is enhanced in sporadic ALS

Given these findings, we next examined cofilin phosphorylation in SALS human lumbar spinal cords compared to controls (**Table 1**), using a specific antibody that detects phosphorylation at Ser3. Western blotting revealed significantly more phosphorylated cofilin in SALS patients compared to controls (3.5-fold, *\*\*p*=0.0034) **(Fig 5A-B)**, in the absence of differences in total cofilin (1.025-fold, *ns p*=0.8936) or loading (GAPDH) **(Fig 5C),** demonstrating that phospho-cofilin is specifically altered. Hence enhanced phosphorylation of LIMK1/2 correlates with more phosphorylation of cofilin in SALS.

**Figure 5.**
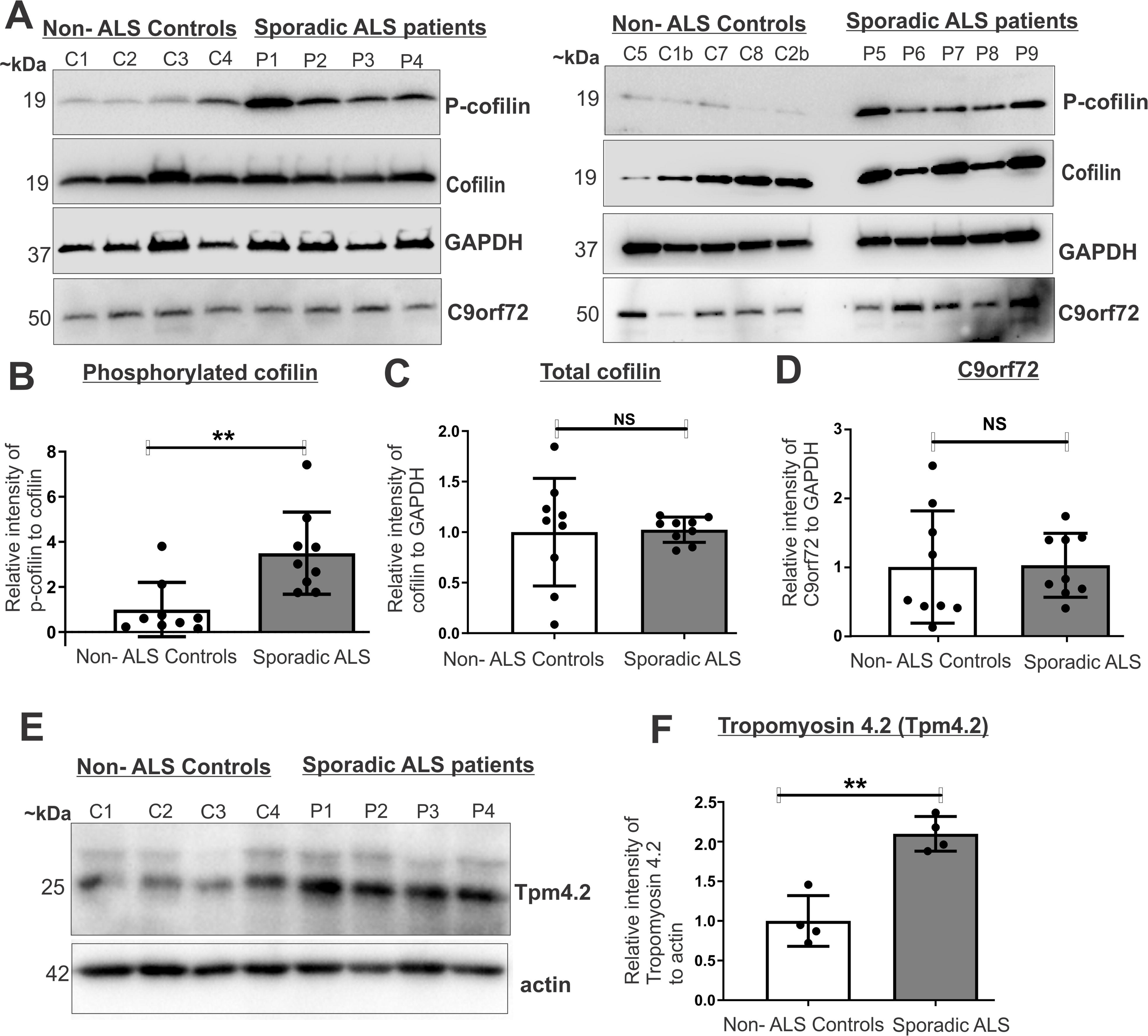
Cofilin phosphorylation and tropomyosin 4.2 are enhanced in sporadic ALS patients. **(A)** Western blotting using anti-phospho-cofilin (Ser3), anti-cofilin, anti-C9orf72 and anti-GAPDH antibodies in lumbar spinal cord lysates prepared from human SALS patients (*n*=9) and non-ALS controls (*n*=7 patients, with two patient samples on both gels, C1=C1b, C2=C2b). Approximate molecular weights (in kDa) are shown on the left. GAPDH was used as a loading control. Blots were reprobed from the same membrane, so loading is equivalent. **(B)** Quantification of western blots shown in (A). Relative intensity of phosphorylated cofilin to total cofilin in spinal cords of SALS patients, normalized to non-ALS controls. Data are represented as mean ± SD, unpaired t-test, two-tailed *\*\*p*<0.01. A significant increase in the phosphorylation of cofilin in human sporadic ALS patients compared to non-ALS controls was detected. **(C)** Quantification of total cofilin relative to loading control GAPDH, and **(D)** Quantification of C9orf72 to relative to GAPDH, both from western blots in (A). No significant differences in levels of cofilin or C9orf72 relative to GAPDH compared to controls was observed, *NS*, non-significant. **(E)** Western blotting using anti-Tpm4.2 and anti-β-actin antibodies of lumbar spinal cord lysates prepared from human SALS patients (*n*=4) and non-neurological controls (*n*=4). Approximate molecular weight markers (in kDa) are shown on the left. Actin was used as a loading control. **(F)** Quantification of western blots shown in (E). The graph depicts the relative intensity of tropomyosin to actin in spinal cords of non-neurological controls and SALS patients. Data are represented as mean ± SD (*n* = 4), unpaired t-test, two-tailed *\*\*p*<0.01. A significant increase in the levels of tropomyosin in human SALS patients compared to non-ALS controls was detected.

Previously, enhanced cofilin phosphorylation was linked to loss of C9ORF72 via haploinsufficiency^53^. Given that 5%–10% of SALS possess hexanucleotide repeat expansions in C9ORF72, we wanted to exclude the possibility that the observed dysregulation of actin dynamics in SALS was due to loss of C9ORF72. Western blotting of SALS lysates confirmed that there were no differences in C9ORF72 levels (1.026-fold, *ns p*=0.9347) **(Fig 5D).** Hence, dysregulation of actin dynamics and aberrant cofilin phosphorylated in SALS is not due to C9orf72 haploinsufficiency.

### Tropomyosin 4.2 is increased in sporadic ALS

Given that Tpm4.2 facilitates cofilin phosphorylation^62,63^, we next examined the SALS lysates by Western blotting using an antibody specific for the Tpm-4.2 isoform. These analyses revealed significantly more Tpm-4.2 in SALS patient lumbar spinal cords compared to controls (2.09-fold, \*\**p*=0.0013) (**Fig 5E-F**). Hence increased levels of this tropomyosin isoform are present in ALS, consistent with the presence of more cofilin phosphorylation and more polymeric actin.

### Cofilin phosphorylation correlates with disease progression in TDP-43 rNLS mice

We next examined whether aberrant cofilin phosphorylation correlates with TDP-43 pathology during disease course **(Supplementary** Fig 1B) in the TDP-43 rNLS mouse model. In these animals, cytoplasmic human TDP-43 accumulates at one week off Dox. At two weeks off Dox, TDP-43 pathology and mild motor impairment first appears. At four weeks, there is major cortical atrophy and muscle NMJ denervation, and by six weeks, there is loss of spinal cord motor neurons and a severe motor phenotype. Surprisingly, these mice undergo partial ‘recovery’ when Dox is returned, manifesting as loss of TDP-43 pathology, prevention of motor neuron loss, and rescue of motor impairment.

Cofilin phosphorylation was examined in cortical lysates during disease course from TDP-43 rNLS and age-matched littermate controls using western blotting. There were no significant differences in cofilin phosphorylation (0.99-fold, *p*=0.9815) **(Fig 6A-B)** nor total cofilin (0.89-fold md, *p*=0.245) **(Fig 6A, 6C),** in the absence of TDP-43 pathology in these mice, one week off Dox. However, at two weeks off Dox, significantly more phosphorylated cofilin (2.1-fold, *\*\*\*p*=0.0008) **(Fig 6D-E)** was present in TDP-43 rNLS mice compared to controls. However, there were no differences in total cofilin (0.998-fold, *ns p*=0.9587) **(Fig 6D, 6F),** so this was not due to increased cofilin expression. Hence, aberrant phosphorylation of cofilin correlates with appearance of TDP-43 pathology. Similarly, at both four weeks (1.623-fold, \**p*=0.0421) and six weeks (1.56-fold \**p*=0.0308) off Dox, significantly more phosphorylated cofilin was present in rNLS mice compared to controls **(Fig 6G, H, J, K),** correlating with cortical atrophy, muscle NMJ denervation and motor impairment. Similarly, no changes in total cofilin were detected at these timepoints (0.959-fold, *p=*0.766) **(Fig 6G, 6I)** (1.03-fold, *p*=0.4477) **(Fig 6J, 6L**). Finally, western blotting of lysates from mice at the recovery phase revealed no significant differences to controls (0.937-fold, *p=*0.7636) **(Fig 6M-N),** and no differences in total cofilin (0.8988-fold, *p*=0.6942) **(Fig 6M, 6O)**. Hence phosphorylation of cofilin correlates with disease and TDP-43 pathology in an ALS mouse model^55^.

**Figure 6.**
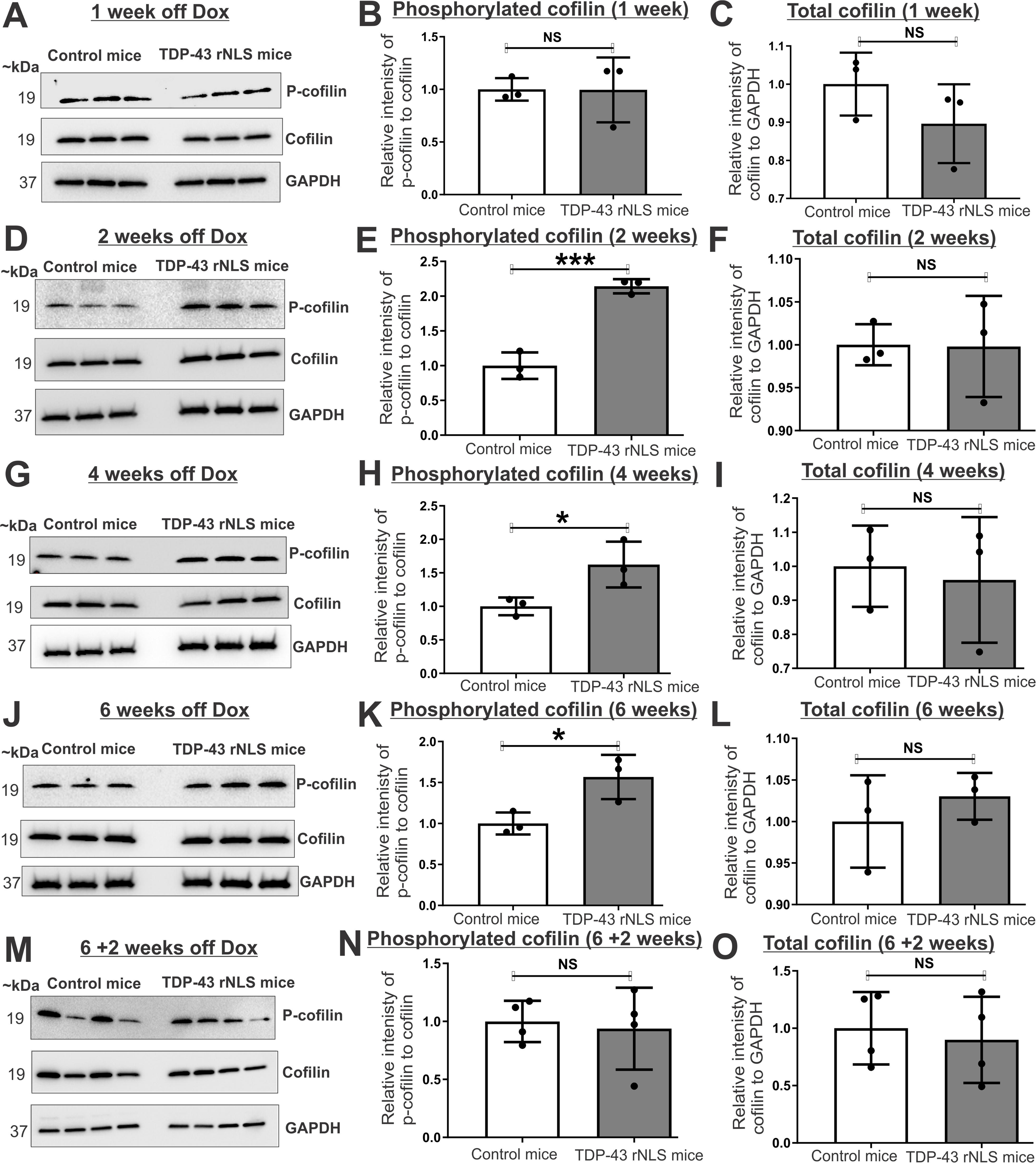
Cofilin phosphorylation increases with disease progression in TDP-43 rNLS mice. Western blotting using anti-phospho-cofilin, anti-cofilin and anti-GAPDH antibodies in lysates prepared from the cortex of littermate control and TDP-43 rNLS mice (*n*=3/group) after 1 week off Dox **(A)**, 2 weeks off Dox **(D)**, 4 weeks off Dox **(G)**, 6 weeks off Dox **(J)** and recovery stage (6 weeks off Dox +2 weeks on Dox) **(M).** The approximate molecular weight (kDa) is shown on the left. GAPDH was used as a loading control**. (B**) Quantification of the western blots in (A). Relative intensity of phosphorylated cofilin relative to total cofilin normalized to control mice after 1 week off Dox. Data are represented as mean ± SD (*n* = 3), unpaired t-test, two-tailed *NS*, non-significant. No significant differences were present in 1-week off DoxTDP-43 rNLS mice relative to control mice. **(E)** Quantification of western blot in (D), mean ± SD (*n* = 3), \*\*\**p*<0.001. **(H)** Quantification of western blot in (G), mean ± SD (*n* = 3), *\*p*<0.05. **(K)** Quantification of western blot in (J), mean ± SD (*n* = 3), *\*p*<0.05. **(E, H, K)** Cofilin phosphorylation is significantly increased in 2, 4 and 6 weeks off Dox TDP-43 rNLS mice compared to control mice**. (N)** Quantification of western blots in (M), mean ± SD (*n* = 3), *NS*, non-significant. No significant difference in phosphorylated cofilin in 6 weeks off Dox + 2 weeks on Dox TDP-43 rNLS mice compared to control mice was present. Quantification of total cofilin levels relative to GAPDH, normalised to controls in 1 week off Dox **(C)**, 2 weeks off Dox **(F)**, 4 weeks off Dox **(I)**, 6 weeks off Dox **(L)** and recovery stage (6 weeks off Dox +2 weeks on Dox) **(O),** mean ± SD (*n* = 3) are *NS*, non-significant. **(C, F, I, L, O).** No significant differences in the levels of total cofilin compared to GAPDH were observed. Blots were reprobed so loading is equivalent.

### Phosphorylated cofilin is enhanced in spinal cord neurons of SALS patients

As phosphorylated cofilin was increased in SALS patients and TDP-43 rNLS mice, next we examined whether this was present specifically in motor neurons in human spinal cords. Immunohistochemistry was performed using SMI-32 and phosphorylated cofilin antibodies. Motor neurons were identified by positive staining for SMI-32 and their size/morphology, as the largest cells in the ventral horn of the spinal cord. Quantification revealed that the levels of phosphorylated cofilin were significantly increased in SALS neurons compared to non-neurological controls (\**\*p*=0.0041) **(Fig 7A-B).** Hence more phospho-cofilin is present in motor neurons in human SALS patients.

**Figure 7.**
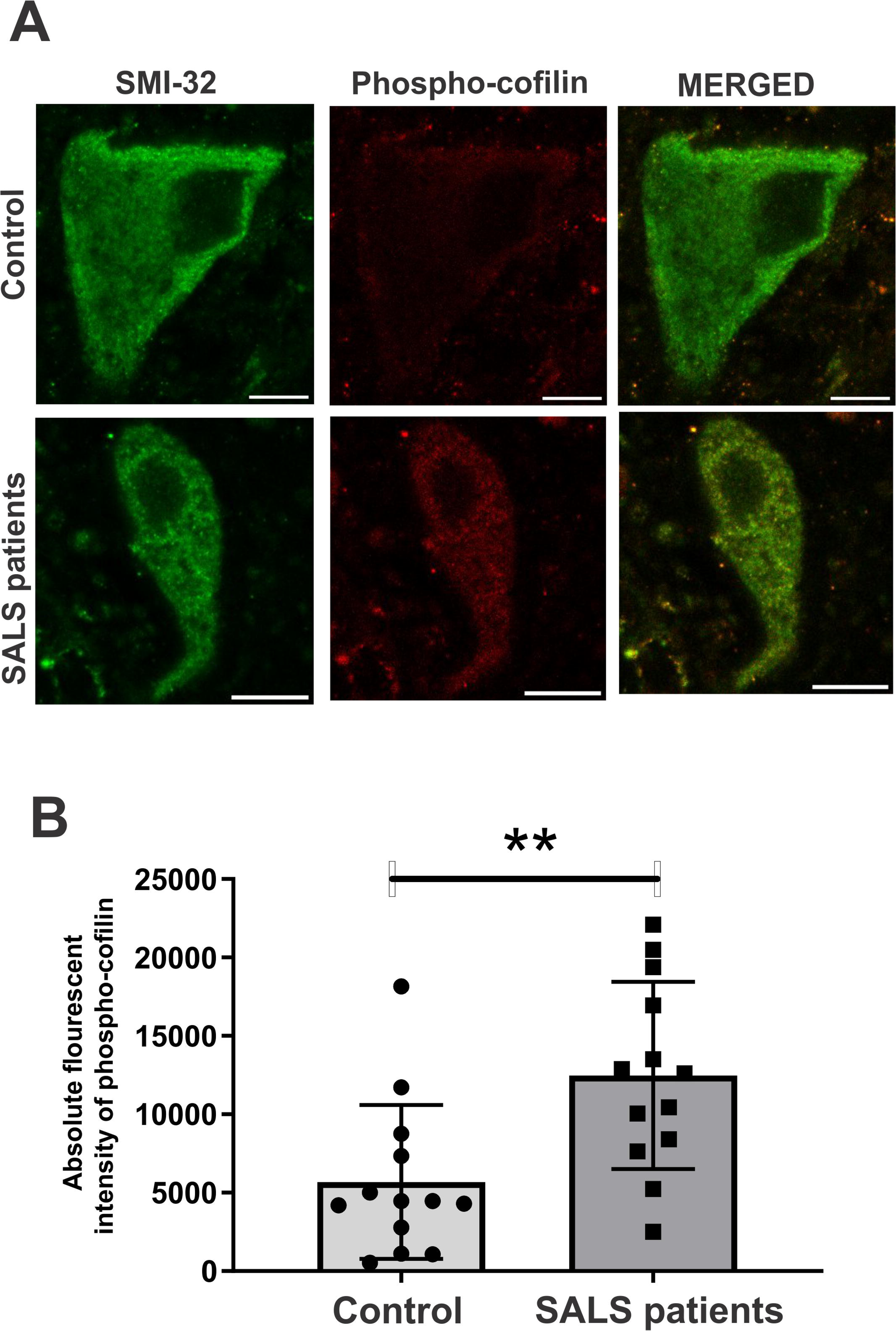
Phosphorylated-cofilin is increased in thoracic spinal cord neurons of SALS patients. **(A)** Immunohistochemistry of post-mortem thoracic spinal cord sections of three sALS patients and two control patients using antibodies against phospho-cofilin and SMI-32. Phosphorylated-cofilin was expressed diffusely in both control patients and in sALS patient motor neurons. Scale bar = 10 μm. (**B)** Quantification of absolute fluorescent intensities revealed increased phospho-cofilin in SALS patients motor neurons compared to non-neurological controls. Data represented from three sALS patients (*n*=13 neurons) and two controls (*n*=13, neurons), unpaired t-test, two tailed, \*\**p*<0.01.

## Discussion

The functions of actin normally depend on its dynamic assembly/disassembly into G-actin and F-actin. Here, we demonstrate that actin dynamics is dysregulated in sporadic ALS. Importantly, this is associated with TDP-43 pathology, the characteristic hallmark present in almost all ALS cases, including SALS^7^ and ∼45% of FTD cases. Moreover, cytoplasmic mis-localisation of TDP-43 is a strong, predictive indicator of toxicity and neuronal death^9^. We detected more actin polymerisation and hence F-actin *in vitro*, in SALS patient lumbar spinal cords, and a mouse model with TDP-43 pathology, that displays key behavioural features of ALS. Furthermore, we detected elevated levels of phospho-cofilin in spinal motor neurons in patients with sporadic disease, demonstrating that these defects are present in the most relevant cell type. These events have two important possible implications in ALS: reduced levels of G-actin and increased levels of F-actin, and an altered G-actin: F-actin ratio, which together will impact the normal functions of actin. This is the first study to correlate TDP-43 pathology to disturbance of actin dynamics and it implies that actin dysregulation is relevant to the pathophysiology of sporadic ALS and FTD.

Actin dynamics is connected to many other cellular process, and of relevance to ALS/FTD, this includes axonal transport,^64–66^ protein quality control,^67–69^ integrated stress responses,^70^ maintenance of cellular redox conditions,^31^ nucleocytoplasmic shuttling,^39^ and DNA damage.^35,71^ Furthermore, the actin cytoskeleton supports a vast number of fundamental processes in neurons, in regulating morphogenesis and structural plasticity, axonal/dendritic initiation, axonal growth, guidance, and branching, and dendritic spine formation and stability.^44,72^ Moreover, actin regulates apoptosis, and possibly other types of cell death described in ALS,^34,73^ including pyroptosis, necroptosis, lysosomal cell death and ferroptosis.^74^ Hence, disturbance of actin dynamics would perturb cellular processes relevant to ALS and thus contribute to neurodegeneration.

We focussed on cofilin as a possible mediator of disrupted actin dynamics. Cofilin is one of the major actin-binding proteins that regulates actin polymerisation.^27^ It functions in many cellular activities and in neurons it regulates neurogenesis, axonal growth, and axonal regeneration.^75^ When phosphorylated, cofilin is inactivated and loses the ability to bind to and depolymerize actin filaments. Here we detected more phosphorylated cofilin in ALS, including in SALS patient motor neurons. More cofilin phosphorylation was previously detected in a small cohort of C9ORF72 ALS/FTD patient samples^53^ and at end stage SOD1^G93A^ mice.^54^ However, we confirmed that increased cofilin phosphorylation in SALS is not due to C9ORF72 haploinsufficiency. Hence, these findings imply that aberrant cofilin phosphorylation may be a common mechanism in different forms of ALS.

We also detected increased LIMK1/2 phosphorylation in SALS patients. This would also enhance cofilin phosphorylation and thus contribute to production of more F-actin. Similarly, Tpm-4.2 also regulates cofilin ^24,25^ and we detected more in SALS, correlating with both increased actin polymerisation and cofilin phosphorylation. Tpm-4.2 interacts and co-polymerises with actin filaments, leading to their stabilisation in both muscle and non-muscle cells.^18^ Thus, it prevents depolymerisation of filaments and enhances actin polymerisation.^17,62,63^ Hence together cofilin, LIMK1/2 and Tpm4.2 likely contribute to actin dysregulation in ALS.

Increased phosphorylation of cofilin was detected early, at symptom onset, and at symptomatic disease stages, in rNLS TDP-43 mice compared to controls. In this model, TDP-43 pathology is present at these timepoints. However, there were no differences in cofilin phosphorylation at the pre-symptomatic disease stage, when mis-localisation of TDP-43 to the cytoplasm is incomplete and post-translational modifications characteristic of pathological TDP-43 (truncation, phosphorylation) are not present.^55,76^ Similarly, no differences in cofilin phosphorylation were evident at the recovery phase, when TDP-43 pathology is absent in this model. Hence these data imply that phosphorylation of cofilin correlates with human TDP-43 pathology and cytoplasmic expression of TDP-43.

We also detected enrichment of insoluble actin in cells expressing TDP-43 ΔNLS, further connecting actin dynamics to TDP-43 pathology. This link was strengthened by the finding that inducing actin polymerisation pharmacologically with Jasplakinolide produced features of TDP-43 pathology (cytoplasmic mis-localisation and inclusion formation).^9,77–79^ This also implies that dysregulating actin dynamics by enhancing actin polymerisation directly induces TDP-43 pathology. Cytoskeletal disruption can lead to abnormal protein-protein interactions, protein misfolding and aggregation,^69,80–83^ implying that disrupted actin dynamics would trigger TDP-43 pathology in ALS. However, it is also possible that TDP-43 pathology directly impairs actin dynamics, so the directionality of these events cannot be confirmed. Alternatively, it is possible that these two mechanisms co-exist and work synergistically.

We also found that Jasplakinolide induced the formation of SGs containing TDP-43. SGs are cytoplasmic ribonucleoprotein structures that form when translation initiation is stalled during cellular stress. They are thought to be initially protective, but during chronic stress they become persistent and may act as a seeding mechanism for the aggregation of TDP-43 and other proteins.^61,84^ Moreover, they are increasingly associated with pathogenesis in ALS and other neurodegenerative diseases.^85^ Our results suggest that actin polymerisation promotes the formation of SGs containing TDP-43 in ALS. Consistent with this notion, actin polymerisation is known to stall transcription and translation during the integrated stress response (ISR),^70,86,87^ a central signalling network that sensors specific types of stress via four different kinases.^70^ The ISR is regulated by phosphorylation of eukaryotic initiation factor 2α (eIF2α) which in turn is modulated by protein phosphatase 1 subunit complex (PPP1R15). G-actin associates with the catalytic subunit of PPP1R15 and reduced G-actin levels induce and fine tune the ISR^70^. Given our finding that G-actin levels were reduced in ALS, it is possible that this activates the ISR and induces cellular stress. One of major processes that induce SG formation is oxidative stress, which has long been implicated in ALS pathophysiology.^79^ Interestingly, actin is one of the most oxidatively modified proteins following oxidative stress ^88,89^ and nitrated actin is significantly increased in peripheral blood mononuclear cells of ALS patients and SOD1^G93A^ rat models, linking oxidative modifications of actin to ALS.^90^ The Hsp104-dependent separation of protein aggregates requires actin filaments as a scaffold for tethering large aggregates,^67,68^ revealing an association between actin, spatial protein quality control and aggregation. Similarly, cytoskeletal motor proteins (kinesin, dynein) also promote TDP-43 SG formation.^91^

TDP-43 normally shuttles between the nucleus and cytoplasm through pores in the nuclear membrane,^57^ but pathological TDP-43 disrupts nucleocytoplasmic shuttling of proteins and mRNA^92^ through cytoplasmic aggregation and mis-localisation of nucleoporins and transcription factors.^92^ This perturbs the morphology of the nuclear membrane and nuclear pore complexes (NPCs),^92^ and defects in nucleocytoplasmic transport are now emerging as an important disease mechanism in ALS/FTD.^1^ Actin plays an important role in maintaining the architecture of the nuclear membrane,^93–95^ and disruption to actin affects the integrity of NPCs.^95^ Likewise, SG formation also disturbs nucleocytoplasmic transport and induces neurodegeneration in ALS.^96^ It is therefore possible that actin dysregulation contributes to nucleocytoplasmic shuttling defects in ALS, although this was not examined here. However, increased actin polymerisation induced by Jasplakinolide led to mislocalisation of TDP-43 to the cytoplasm, demonstrating that aberrant nucleocytoplasmic shuttling of TDP-43 is associated with dysregulation of actin dynamics in ALS.

Disturbances to actin dynamics have also been described in other neurodegenerative diseases,^91,97–99^ implying that actin dysregulation contributes to the degeneration of neurons. Abnormal cofilin-actin structures (‘rods’) are present in tissues of Alzheimer’s and Parkinson’s disease patients^97,40,91,98^ and plexin–actin rods are pathogenic structures associated with spinal muscular atrophy (SMA).^100^ Interestingly, TDP-43 inclusions have also been detected in over 50% of AD cases,^101^ further highlighting the relationship between aberrant actin dynamics, cofilin and TDP-43 pathology. Furthermore, Hirano bodies are actin-containing inclusions present in Alzheimer’s disease (and other neurodegenerative conditions) that also contain actin-associated proteins, tropomyosin, α-actinin and vinculin.^102,103^ Actin-based structures were recently identified in adult motor neurons from symptomatic SOD1^G93A^ mice, which enhanced neurite outgrowth and branching.^104^

Increased polymerisation of actin in ALS could be mediated by its interaction with TDP-43 because we detected a possible physical association between TDP-43 and actin via immunoprecipitation, using both purified proteins and cell lysates. Inclusions immunoreactive for β-actin have been previously detected in SALS patient tissues, although the protein/s interacting with actin were not identified.^105^ Similarly, actin-containing Hirano bodies are present in ALS, as well as other neurodegenerative conditions.^102,106,107^ TDP-43 interacts with components of microtubules, neurofilaments, and actin cytoskeletal proteins^108^ and cytoplasmic accumulation of TDP-43 *in vitro* and *in vivo* diminishes axonal growth.^109^ Accumulation of both TDP-43 and alpha-actin was previously detected in oligodendrocytes and neurons of familial ALS patients bearing Angiogenin variants.^110^ Similarly, actin interacts with other misfolded proteins associated with neurodegeneration, resulting in accumulation of F-actin. Actin interacts with tau, inducing its abnormal polymerisation into F-actin *in vitro* and *in vivo* in *Drosophila* models of Alzheimer’s disease.^99^ F-actin also interacts with presenilin and tau inclusions,^111,112^ implying that aberrant accumulation of F-actin is instrumental in pathological changes in neurons. TDP-43 is also known to regulate several cytoskeletal RNAs as part of its function in RNA metabolism, including actin.^108^ Together, these findings suggest that actin disturbances are linked to TDP-43 pathology by a physical interaction between actin and TDP-43.

In summary, this study provides novel insights into the pathophysiology of ALS/FTD, revealing that TDP-43 pathology is closely related to dysregulated actin dynamics. Mechanistically this could result from aberrant phosphorylation of cofilin and LIMK1, Tpm-4.2, and/or an interaction between TDP-43 and actin. Given its relationship to other pathways implicated in ALS, dysregulated actin dynamics could be a missing link that precedes SG formation, dysregulation of nucleocytoplasmic transport, and neurodegeneration in ALS/FTD. A hypothetical model illustrating the relationship between TDP-43 and dysregulation of actin in ALS is shown in **Figure 8**. This highlights cofilin as the central player in this complex web of biochemical alterations. Hence strategies to modulate cofilin activity, by inhibiting its phosphorylation and thus increasing its activity, may be a novel therapeutic strategy in ALS, FTD and other diseases in which TDP-43 pathology is a central component.

**Figure 8.**
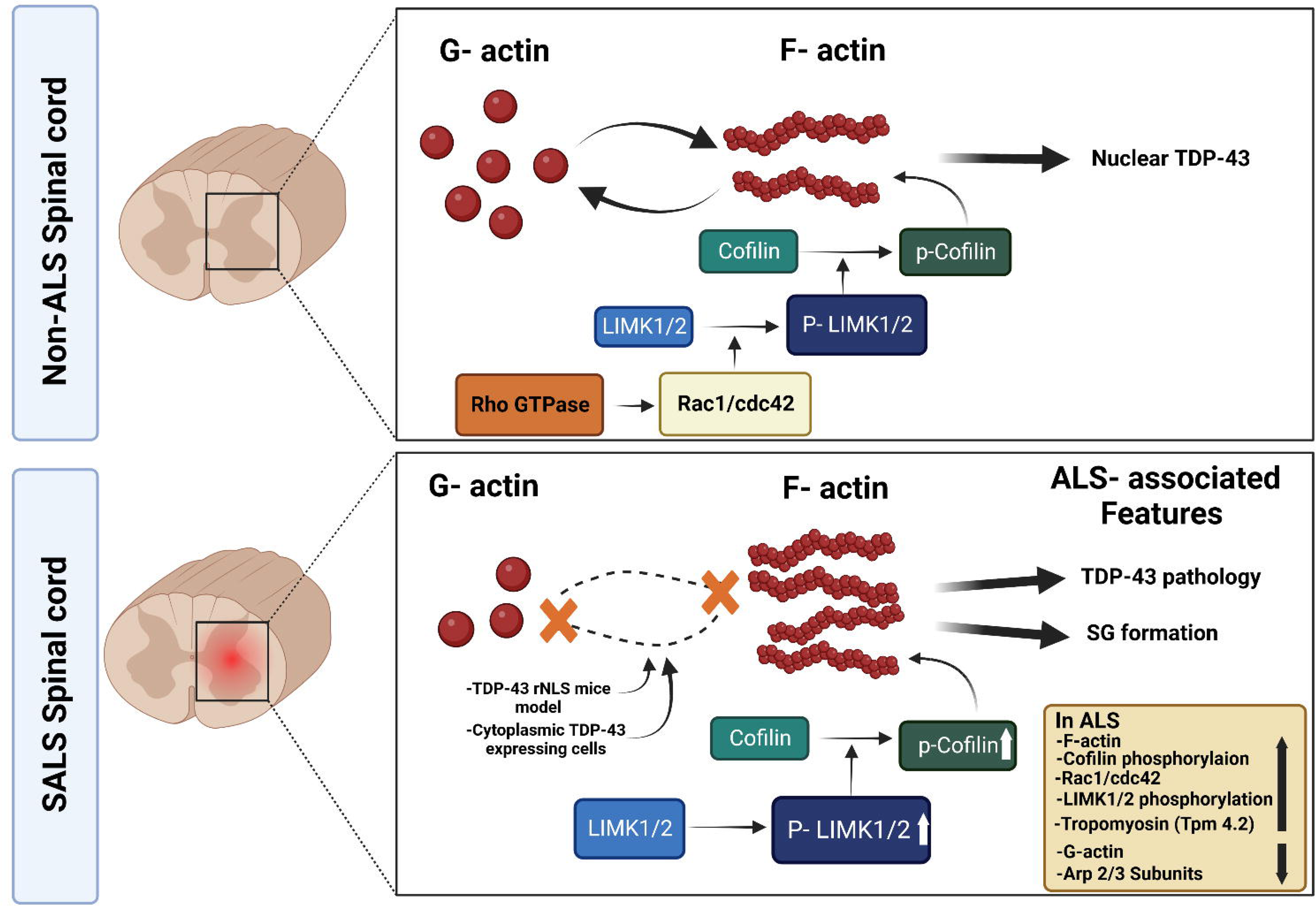
The LIMK1/2-cofilin pathway is dysregulated in sporadic ALS and associated with TDP-43 pathology. A hypothetical model illustrating the relationship between TDP-43 pathology and actin dynamics in normal physiology (*top panel*) and in ALS (*bottom panel*). Rac1/cdc42, Tpm 4.2 and phosphorylated forms of both cofilin and LIMK1/2 are increased in sporadic ALS (SALS) patients. This leads to abnormal actin dynamics and increased formation of filamentous F-actin. This promotes the production of pathological forms of TDP-43, involving its cytoplasmic mis-localisation, aggregation, and stress granule (SG) formation. Hence, this study identifies dysregulation of actin dynamics as a disease mechanism associated with TDP-43 pathology in sporadic ALS.

## Acknowledgements

The authors thank Dr. Adam Walker, University of Queensland for providing constructs and TDP-43 rNLS mice tissue lysates, and Dr. Damian Spencer and Prof. Begona Heras, La Trobe University for providing LICA-TDP43 constructs.

## Ethics statement

This study was approved by Macquarie University Human Ethics (HEC # 5201600387 and #5201700825) and Macquarie University Animal Ethics Committees (ARA # 2015/042).

## Supplementary Material

All full-length blots are shown in the supplementary material. Supplementary material is available at *Brain* online.

