## Supplemental figures for "Dysregulated actin dynamics and cofilin correlate with TDP-43 pathology in sporadic amyotrophic lateral sclerosis"

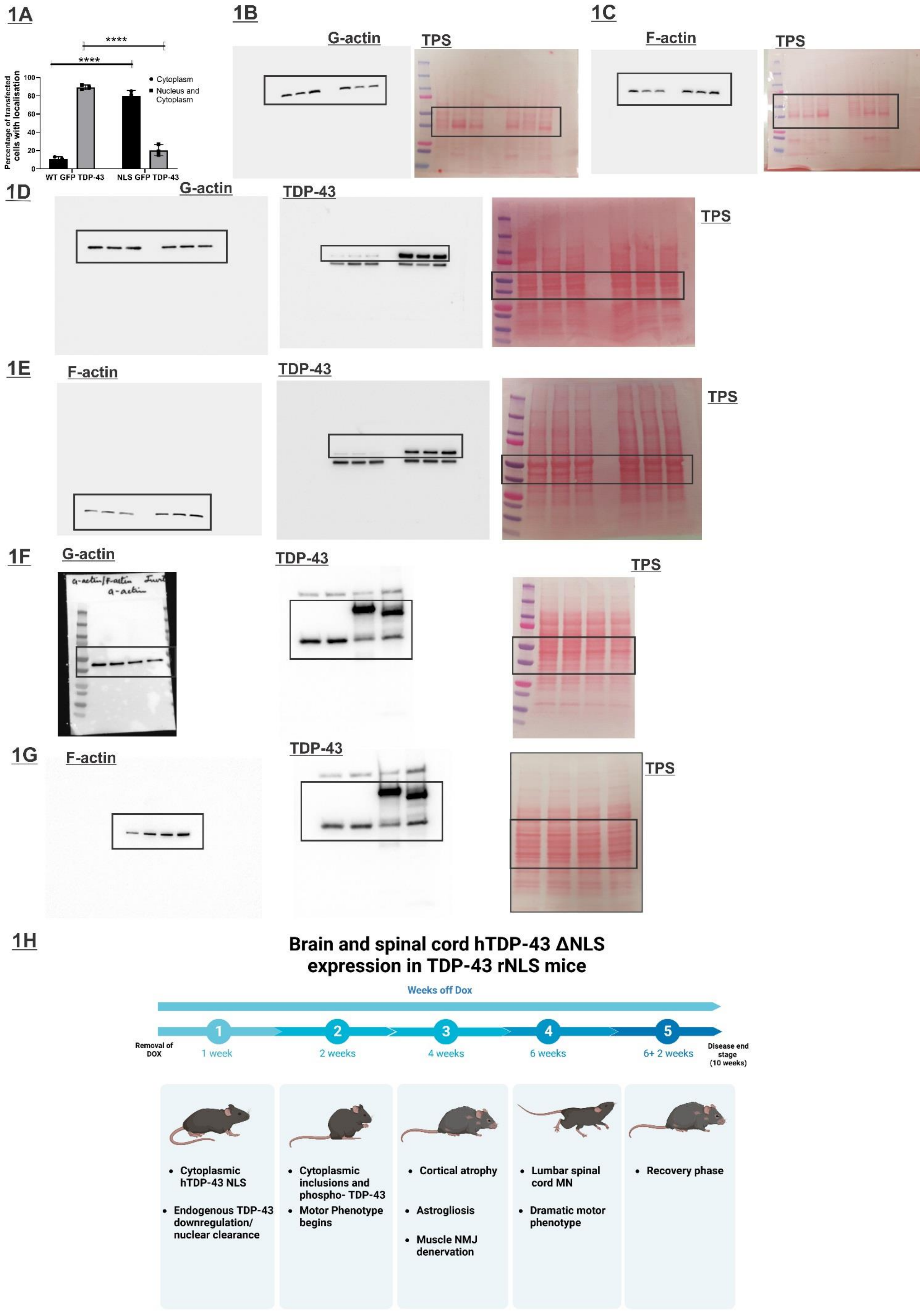
1A
****
****
Percentage of transfected cells with localisation
100
80
60
40
20
0
Cytoplasm
Nucleus and Cytoplasm
WT GFP TDP-43
NLS GFP TDP-43
1B
G-actin
TPS
1C
F-actin
TPS
1D
G-actin
TDP-43
TPS
1E
F-actin
TDP-43
TPS
1F
G-actin
TDP-43
TPS
1G
F-actin
TDP-43
TPS
1H
Brain and spinal cord hTDP-43 ΔNLS expression in TDP-43 rNLS mice
Weeks off Dox
Removal of DOX
1 week
2 weeks
4 weeks
6 weeks
6+ 2 weeks
Disease end stage (10 weeks)
Cytoplasmic hTDP-43 NLS
Endogenous TDP-43 downregulation/ nuclear clearance
Cytoplasmic inclusions and phospho- TDP-43
Motor Phenotype begins
Cortical atrophy
Astrogliosis
Muscle NMJ denervation
Lumbar spinal cord MN
Dramatic motor phenotype
Recovery phase

**<u>Supplementary Figure 1</u>**

**(1A) Localisation of TDP-43 expressed in NSC-34 cells** Quantification of transfected NSC-34 cells expressing wild-type TDP-43 GFP or TDP-43ΔNLS GFP based on the localisation of TDP-43. Cells were categorised as either cytoplasmic only or as both nuclear and cytoplasmic, which included cells with TDP-43 localised in both the nucleus and cytoplasm and those with nuclear TDP-43 only. A total of >100 cells per group were examined. Data are represented as mean ±SD, n=3, two-way Anova, Sidak's multiple comparison test ****p<0.001.

(**1B and 1C) Full-length blots of Figures 1A and 1C.**

**(1D and 1E) Full- length blots of Figures 1F and 1H.**

**(1F and 1G) Full- length blots of Figures 1K and 1M**

**(1H) Timeline of disease stages in TDP-43 rNLS mice.** Cytoplasmic expression of TDP-43 begins at one week off Dox but is incomplete at this stage **(pre-symptomatic stage**). Cytoplasmic inclusions and phospho-TDP43 are detected in the cortex of these mice at two weeks off Dox, and a motor phenotype begins at this stage **(symptomatic onset stage)**. At four weeks off Dox, mice display decreased cortical thickness accompanied by astrogliosis, indicating neurodegeneration **(early disease stage)**. A dramatic motor phenotype and loss of 30% of lumbar spinal cord motor neurons occur at 6 weeks off Dox **(late symptomatic stage).** Phospho-TDP-43 inclusions are also detected in the motor cortex at this stage. Re-introduction of Dox for two weeks eliminates cytoplasmic phosphorylated TDP-43, leading to decreased cortical atrophy and diminished astrogliosis **(recovery stage).**

## 2A

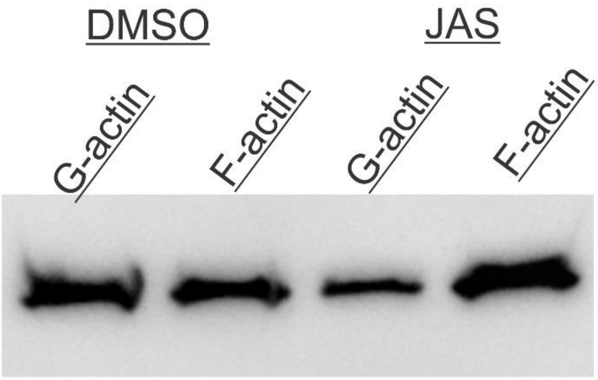


## 2B

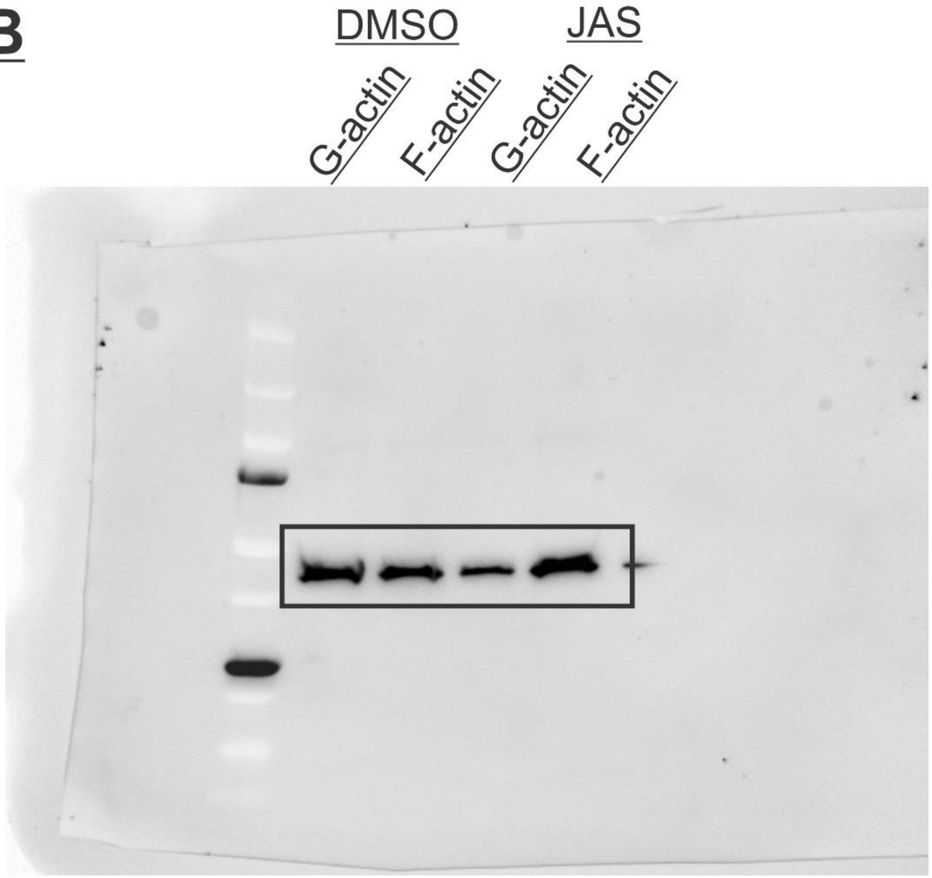


<u>**Supplementary Figure 2**</u>

**(A) Jasplakinolide induces the production of F-actin.** Western blotting of G-actin and F-actin fractions from NSC-34 cells after treatment with DMSO or 1µM Jasplakinolide (JAS). Actin polymerisation is increased in NSC-34 cells treated with 1µM Jasplakinolide**.**
**(B) Full- length blot shown in Supplementary Figure 2(A)**

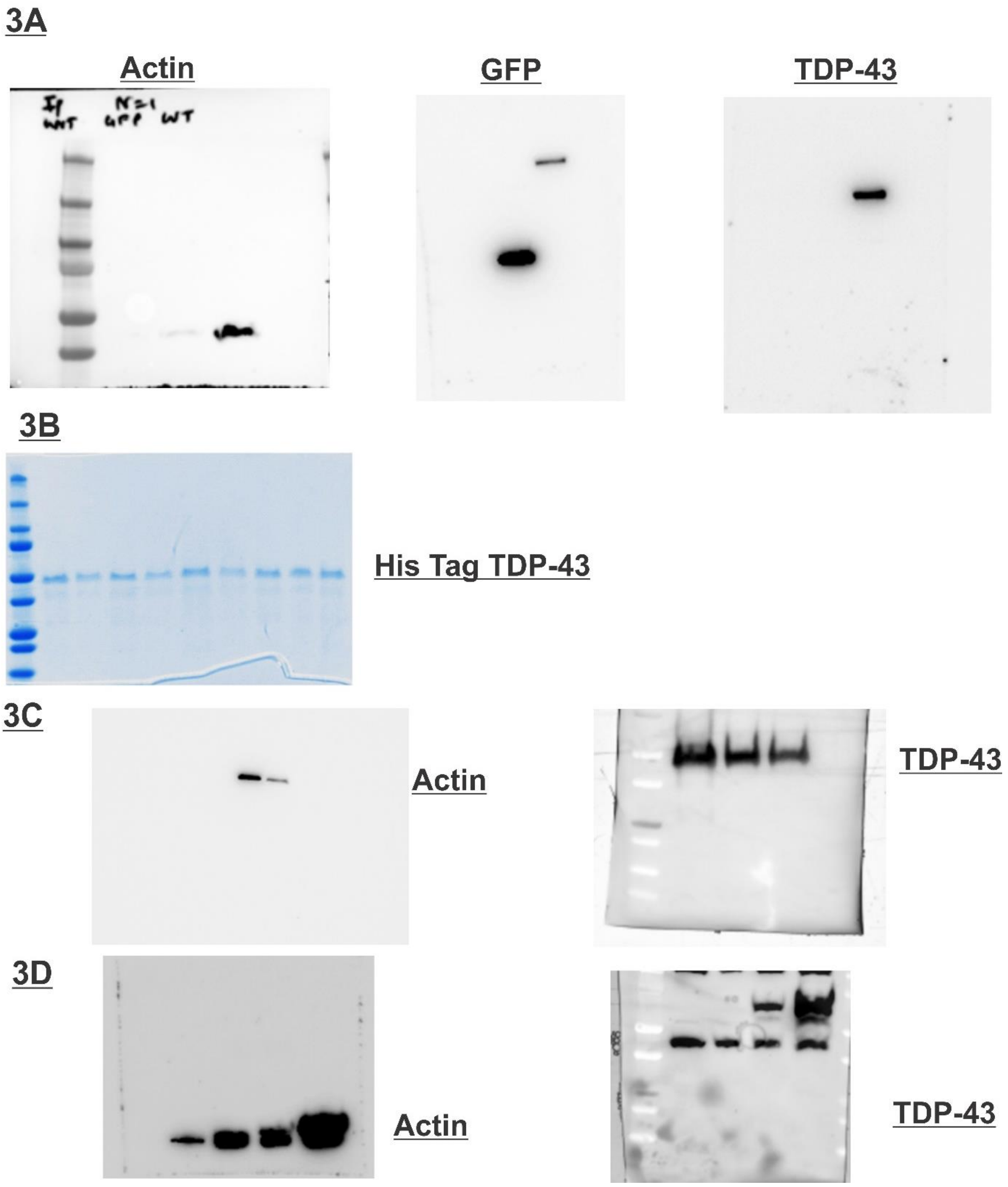


**Supplementary Figure 3**

**(A)** **Full-length blots of Figure 3A.**

**(B)** **Purified recombinant TDP-43 protein** Coomassie staining of purified TDP-43 WT following SDS-PAGE. Molecular weight markers are shown on the left. E1-E9 represent different eluted fractions, each of volume 5$\mu$L.

**(C, D)** **Full-length blots of Figures 3C and 3D.**

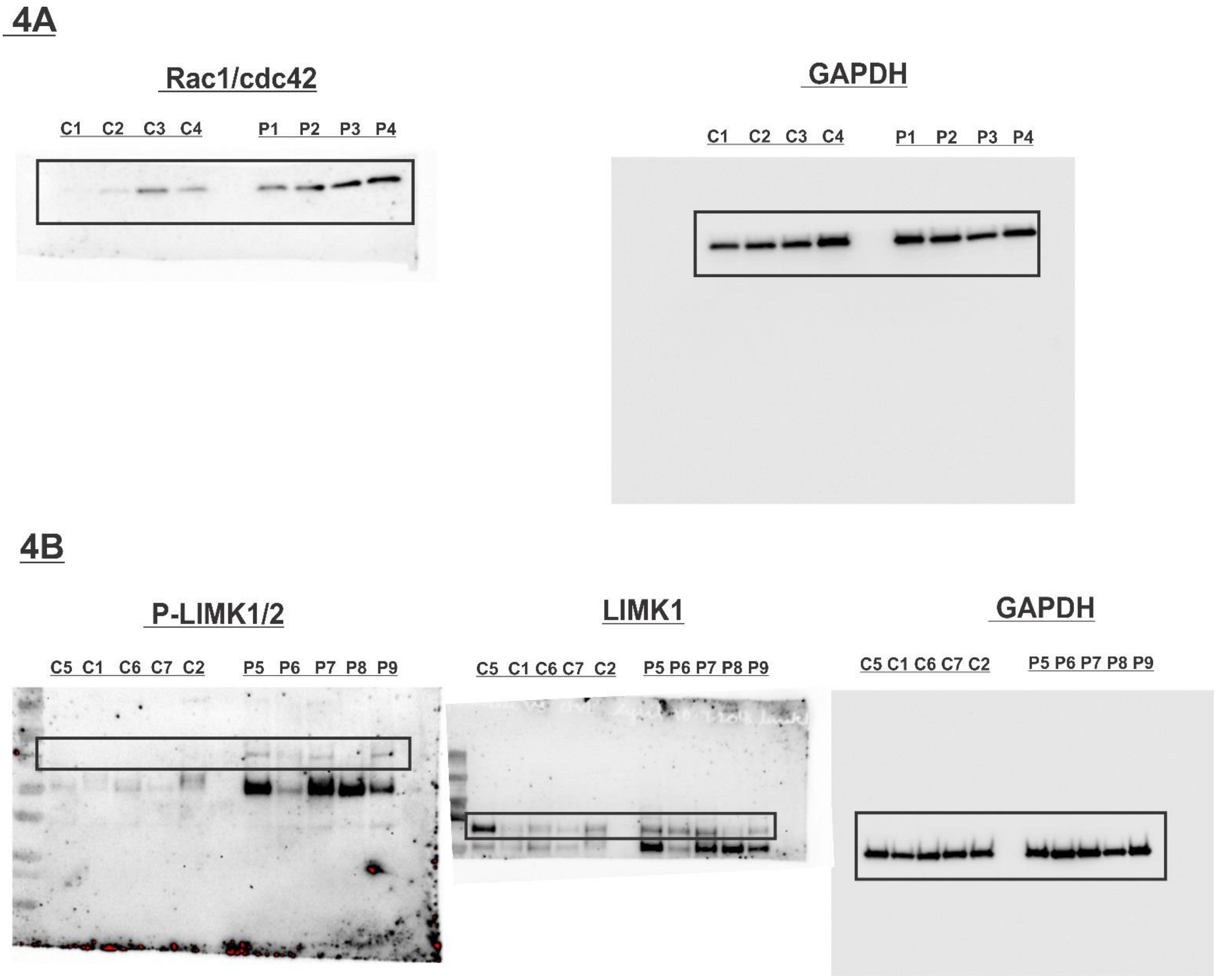


**Supplementary Figure 4**

**(A, B) Full- length blots corresponding to Figures 4A and 4C.**

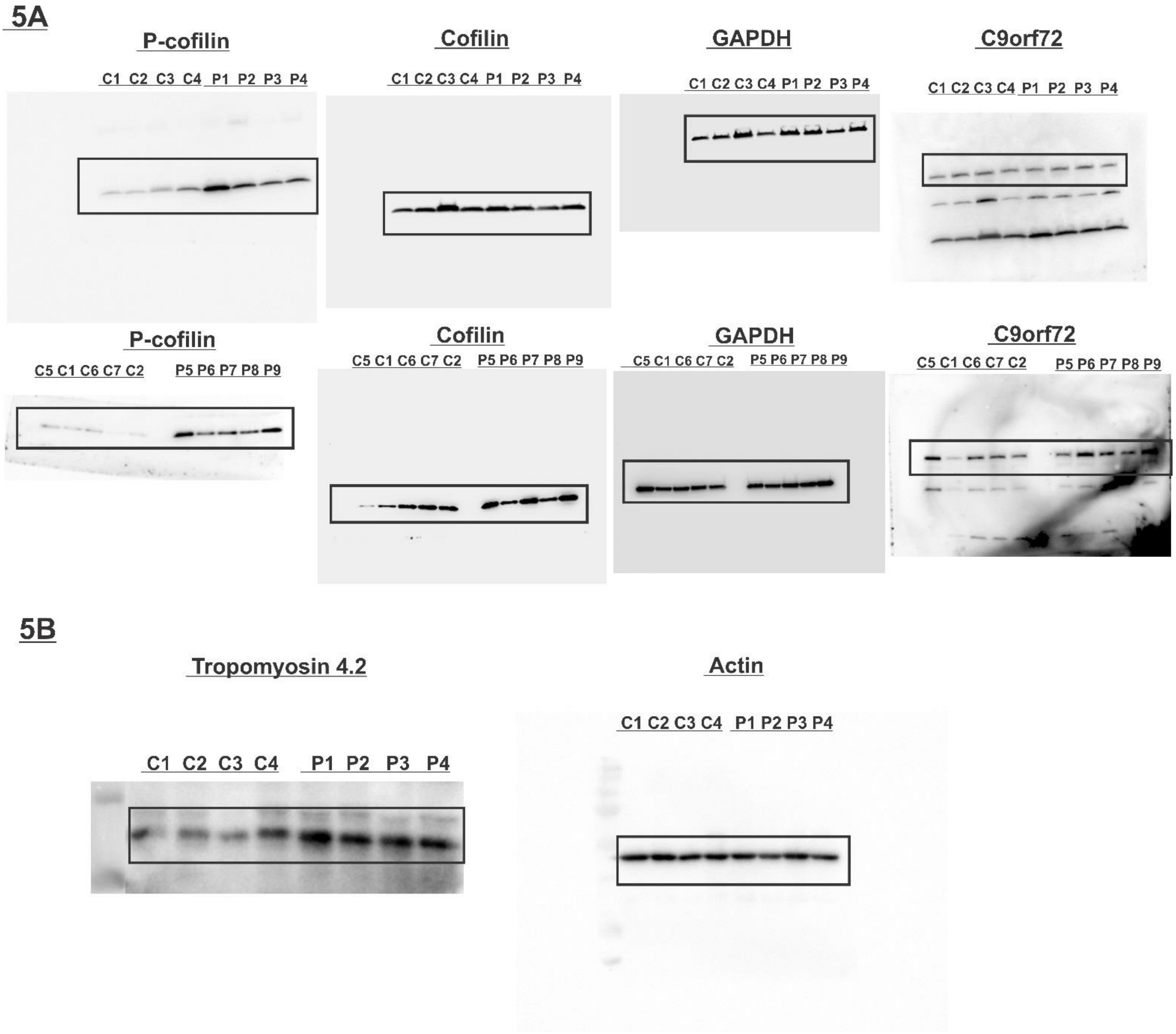


**Supplementary Figure 5**

**(A, B) Full- length blots corresponding to Figures 5A and 5E.**

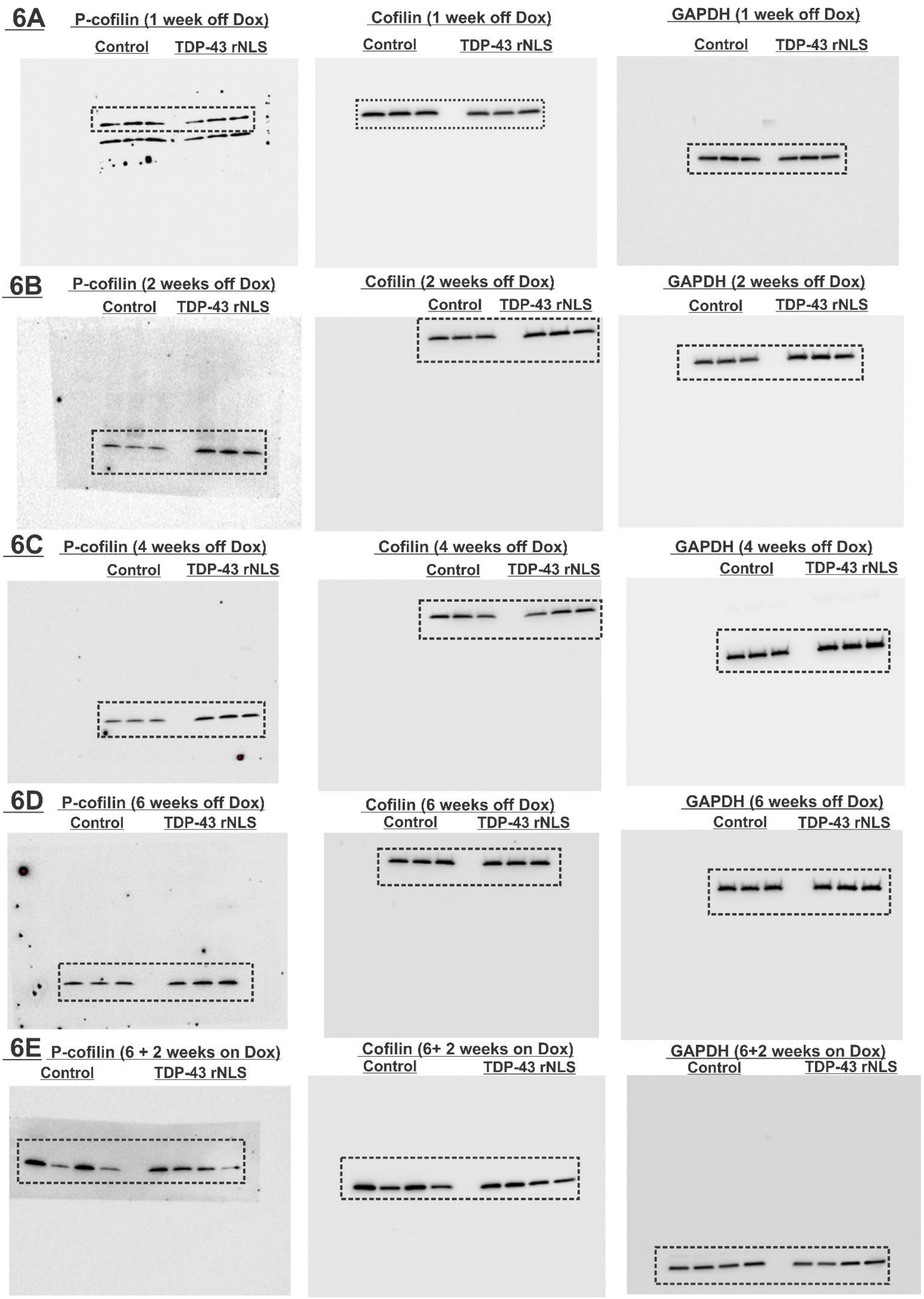
6A
P-cofilin (1 week off Dox)
Control
TDP-43 rNLS
Cofilin (1 week off Dox)
Control
TDP-43 rNLS
GAPDH (1 week off Dox)
Control
TDP-43 rNLS
6B
P-cofilin (2 weeks off Dox)
Control
TDP-43 rNLS
Cofilin (2 weeks off Dox)
Control
TDP-43 rNLS
GAPDH (2 weeks off Dox)
Control
TDP-43 rNLS
6C
P-cofilin (4 weeks off Dox)
Control
TDP-43 rNLS
Cofilin (4 weeks off Dox)
Control
TDP-43 rNLS
GAPDH (4 weeks off Dox)
Control
TDP-43 rNLS
6D
P-cofilin (6 weeks off Dox)
Control
TDP-43 rNLS
Cofilin (6 weeks off Dox)
Control
TDP-43 rNLS
GAPDH (6 weeks off Dox)
Control
TDP-43 rNLS
6E
P-cofilin (6 + 2 weeks on Dox)
Control
TDP-43 rNLS
Cofilin (6+ 2 weeks on Dox)
Control
TDP-43 rNLS
GAPDH (6+2 weeks on Dox)
Control
TDP-43 rNLS

**Supplementary Figure 6**

**(A, B) Full- length blots corresponding to Figures 6A and 6D.**

**(C, D, E) Full-length blots corresponding to Figures 6G, 6J and 6M.**
